# Emetine dihydrochloride inhibits Chikungunya virus nsP2 helicase and shows antiviral activity in the cell culture and mouse model of virus infection

**DOI:** 10.1101/2025.06.20.660672

**Authors:** Anshula Sharma, Chandru Subramani, KA Shouri, Ghanshyam Sharma, Brohmomoy Basu, Abhay Deep Pandey, Archana Rout, Devendra Sharma, Deepti Jain, Sudhanshu Vrati

## Abstract

Chikungunya virus (CHIKV), a mosquito-borne alphavirus, causes frequent epidemics of chikungunya fever across the world and has become a significant medical challenge, warranting the development of novel antivirals. A cell imaging-based high-content screening of the Spectrum collection of small molecules, containing all the US and international drug molecules, identified Emetine dihydrochloride (ED) as a potent antiviral against CHIKV. ED inhibited CHIKV replication in the C57BL/6 mouse model of chikungunya disease, resulting in significantly lower viremia. Importantly, clinical symptoms of joint swelling were altogether absent in the CHIKV-infected mice treated with ED. In the ERMS cells, ED inhibited the CHIKV uptake, and the virus replication early during the virus life cycle by inhibiting the viral RNA synthesis, resulting in the inhibition of the viral protein synthesis. ED was predicted to bind the helicase domain of the CHIKV non-structural protein nsP2 by *in silico* methods. The ED binding to the nsP2 protein and its helicase domain was validated *in vitro* by microscale thermophoresis and isothermal titration calorimetry. Importantly, ED inhibited the CHIKV nsP2 helicase activity *in vitro* in a dose-dependent manner. These data demonstrate the potential of ED to be repurposed as a novel CHIKV antiviral, warranting clinical development.

**AUTHOR SUMMARY:** Chikungunya virus (CHIKV) spreads through the bites of the virus-infected *Aedes* mosquitoes. The virus causes frequent epidemics of chikungunya fever involving severe joint edema and muscle pain. The virus activity has been recorded in more than 100 countries in Asia, Africa, South and Central America, and Europe, and it has become endemic in several regions of the world. In 2024, around 480,000 CHIKV cases and over 200 deaths were reported worldwide. While a CHIKV vaccine was recently approved, no virus-specific antiviral therapy is available. Considering the global medical significance of the virus, the development of effective CHIKV antivirals is a priority. With a view to repurposing an existing drug, we screened a collection of all the approved US and international drug molecules and identified Emetine dihydrochloride (ED) as a potent inhibitor of CHIKV replication in the cell culture model. The CHIKV-infected mice treated with ED showed reduced virus replication and no joint swelling, which is a typical clinical symptom of CHIKV disease. Our studies show that ED binds the CHIKV nsP2, inhibiting its helicase action, which is required for replicating the viral genomic RNA and producing the viral proteins. In the past, emetine has been used in humans to treat amoebiasis. Our work thus shows the potential of the drug emetine to be repurposed for treating CHIKV patients.

## INTRODUCTION

Chikungunya virus (CHIKV) is an arthropod-borne alphavirus within the *Togaviridae* family. The virus is primarily transmitted through *Aedes aegypti* and *Aedes albopictus* mosquitoes. First identified during a 1952–1953 epidemic in Tanzania (1), CHIKV has since caused outbreaks worldwide, including in Africa, Asia, the Americas, Europe, and Australia (2). The notable outbreaks of CHIKV include its periodic epidemics in Africa and Asia from the 1960s to 2000s, a large-scale epidemic in the Indian Ocean region, particularly on Réunion Island in 2005–2006, and a major outbreak in India in 2006 with 1.25 million cases. Rapid CHIKV outbreaks occurred in the Caribbean and South America during 2013–2014, marking its arrival in the Western Hemisphere. From 2016 to 2023, CHIKV has continued to cause frequent outbreaks across Asia, Africa, and the Americas, spreading to over 110 countries worldwide. In 2024 and as of the 30th of November, approximately 480,000 CHIKV cases and over 200 deaths were reported worldwide (3). In addition to urban transmission cycles involving humans, CHIKV engages in a sylvatic transmission cycle with non-human primates serving as amplification hosts and reservoirs during the inter-epidemic periods (1).

CHIKV infection in humans is associated with febrile illness characterised by high fever, maculopapular rash, myalgia, polyarthralgia, and headaches. While many symptoms typically resolve within a week, persistent joint pain may last for months or even years in some patients (4,5). CHIKV infection is typically a self-limiting illness with a low fatality rate of approximately 0.1%. However, the frequent occurrence of joint complications, which often result in persistent disability, poses significant public health challenges. These complications notably affect the quality of life of infected individuals and impose considerable economic and societal burdens, especially in low- and middle-income countries (6). The virus’s potential for vertical transmission, rapid evolution, human travel, and establishment in new geographic regions pose a significant public health challenge (7). These factors underscore the urgent need for effective measures to control CHIKV, including the development of safe and targeted antiviral therapeutics.

CHIKV has an ∼11.8 kb long single-stranded positive-sense RNA genome with a 5’ methyl guanylate cap and a 3’ polyadenylate tail (8). During replication, the viral genome encodes two polyproteins, namely, non-structural and structural polyproteins. The non-structural polyprotein is processed into four non-structural proteins (nsP1, nsP2, nsP3, and nsP4), while the structural polyprotein is cleaved to give rise to five structural proteins (capsid, E3, E2, 6K, and E1). The virus relies on non-structural proteins for genome replication as nsP1 caps viral RNA, nsP2 processes polyproteins by its protease activity, and unwinds replication intermediate dsRNA by its helicase activity, nsP3 aids replication complex formation, and nsP4 acts as the RNA-dependent RNA polymerase. Structural proteins facilitate assembly and infection; the capsid protein encases viral RNA, E1 enables membrane fusion, E2 mediates host receptor binding, E3 aids glycoprotein processing, and 6K supports glycoprotein trafficking and virion assembly (9,10). These proteins collectively drive CHIKV replication, assembly, and release, enabling the virus to propagate within the host.

Several viral proteins have been investigated as potential targets for direct-acting antiviral development (11–16). Various host-targeting antivirals have also been identified by targeting specific host proteins involved directly in the virus replication process (17–21). With the advent of robust cell-based high-throughput screening methods, libraries of a large number of drug-like small molecules, compounds from chemical libraries, and natural sources have been screened for their potential anti-CHIKV activity (15,16,22–28). Virtual screening and computational methods have also been employed to identify potential CHIKV inhibitors (29–34). However, most candidates require further validation in animal models and clinical trials. Few natural products have also been identified as potential antiviral agents (15,33). Despite these efforts, no antiviral therapies have advanced to clinical use, emphasising the need for continued research to develop effective CHIKV interventions for at-risk populations.

In this study, we performed a cell-based high-throughput screening of 2560 compounds from the Spectrum Collection of small molecules that includes all the compounds in the US and international drug collections. We identified four potential antiviral compounds, Niclosamide, Gambogic acid, Celastrol, and Emetine dihydrochloride (ED), that were further evaluated for their antiviral efficacy in the animal model. Among these, ED emerged as the most potent CHIKV inhibitor, exhibiting low cytotoxicity and significant antiviral activity at nanomolar concentrations in the cell culture. Moreover, ED significantly inhibited viremia and alleviated CHIKV-associated joint swelling symptoms in the mouse model of CHIKV infection. Investigations on ED’s antiviral mechanism revealed that it strongly bound to the CHIKV non-structural protein nsP2, inhibiting its RNA helicase activity, and thereby suppressing the viral replication.

## MATERIAL AND METHODS

### Ethics statement

For the animal experiments, the guidelines on the care and use of laboratory animals provided by the Committee for the Purpose of Control and Supervision of Experiments on Animals (CPCSEA), Government of India, were followed. The experimental protocol was approved by the Institutional Animal Ethics Committee of the Regional Centre for Biotechnology (RCB/IAEC/2019/047).

### Cell lines and viruses

BHK-21 (baby hamster kidney cells) and Vero (African green monkey kidney cells) cell lines were obtained from the National Centre for Cell Sciences (India) cell repository. ERMS (human embryonal rhabdomyosarcoma) cells were obtained from the ATCC, USA (RD-CCL-136). BHK-21 were cultured and maintained in 1× Minimum Essential Medium Eagle (MEM) (HiMedia, AL0475) supplemented with 10% Fetal Bovine Serum (FBS) (Gibco), 100 U/ml penicillin and 100 mg/ml streptomycin (HiMedia, A001A). ERMS cells were cultured in Dulbecco Minimum Essential Medium (DMEM) (HiMedia, AL007A) with 10% FBS. The cells were grown in a humidified 5% CO_2_ incubator at 37 °C. The recombinant CHIKV-LR-5’GFP virus strain (15,35) was used for the primary screening of compounds. Further validation involving both *in vitro* and *in vivo* assays was carried out using the wild-type IND-06-Guj isolate of CHIKV (GenBank, JF274082.1). Both viruses were cultured in BHK-21 cells. The virus titer was determined as plaque-forming units per millilitre (pfu/ml) on Vero cells as described before (15).

### Small molecule library

The Spectrum Collection (MicroSource Discovery Systems Inc., USA), containing biologically active and structurally diverse compounds, was used for screening the anti-CHIKV compounds. The collection contained 2560 compounds, including all the US and international drug collections. Besides, several natural and discovery molecules were also included in the collection. The collection was available as a 10 mM DMSO solution of the compounds stored at -80 °C.

### High-throughput drug screening

BHK-21 cells, known for their ability to efficiently replicate CHIKV, ease of culture, high growth rate, and resilience to frequent passages, were utilised for antiviral screening assays. For primary screening, each library compound was tested at a concentration of 10 µM (diluted in DMSO). BHK-21 cells (10,000 cells/well) were seeded in a 96-well black, flat clear-bottom plate (Corning, 3904) in MEM supplemented with 2% FBS and incubated for 24 h at 37 °C. The cell monolayers were treated with 10 µM of each test compound and infected with CHIKV-LR-5’GFP at a multiplicity of infection (MOI) of 0.1. The plates were then incubated for 24 h at 37 °C. 0.1% DMSO, used as the vehicle for diluting the test compounds, was employed as the negative control. The cells were stained with Hoechst 33342 (Thermo Scientific, 62249) and SYTOX Orange Nucleic Acid Stain (Invitrogen, S11368). The plate imaging was performed using the ImageXpress Micro Confocal High Content Imaging System (Molecular Devices) according to the manufacturer’s instructions. Imaging was conducted using DAPI (4′,6-diamidine-2′-phenylindole dihydrochloride) for determining the total cell counts, FITC (Fluorescein isothiocyanate) for CHIKV-infected cells, and Texas Red channels (SYTOX) for dead cells using a 10× Plan Apo NA 0.45 objective. Data analysis was performed with the Multi-Wavelength Cell Scoring Algorithm in fast mode. The antiviral activity of each compound was determined by calculating the percentage inhibition of the GFP fluorescence. Compounds displaying >80% fluorescence inhibition (virus replication inhibition) and >80% viability, with respect to the DMSO vehicle-treated, CHIKV-infected control, were selected as positive hits. The secondary screening involved studying the anti-CHIKV activity of the selected compounds at lower concentrations in ERMS cells.

### Selectivity Index (SI) determination

ERMS cells were used to determine the selected compounds’ 50% CHIKV inhibitory concentration (IC_50_). The cells were seeded at a density of 10^6^ cells/well in the 96-well black flat clear-bottom plates (Corning, 3904) in DMEM supplemented with 2% FBS and incubated for 24 h at 37 °C. The cells were then treated with two-fold serial dilutions of the test compound (50 µl/well) in DMEM with 2% FBS, followed by infection with CHIKV-LR-5’GFP (MOI 5) and incubation for 30 h. The cells were then stained, and the virus replication inhibition was determined using high-content imaging, as described above. To determine the test compound’s 50% cytotoxic concentration (CC_50_), ERMS cells were seeded in 96-well plates (10^6^ cells/well) in DMEM with 2% FBS and incubated at 37 °C for 24 h. The cells were then treated with two-fold serial dilutions of the test compound (50 µl/well) and incubated for 30 h. The cells were stained with DAPI and SYTOX and analysed by high-content imaging. The percentage cell cytotoxicity was calculated from the cell viability in the test compound-treated cells compared to the DMSO-treated vehicle control. All the experiments were performed in triplicate. The IC_50_ and CC_50_ values were determined using the GraphPad Prism software, version 8.0, via nonlinear regression analysis. Each compound’s selectivity index (SI) was calculated as SI = CC_50_/IC_50_. For these experiments, as well as the subsequent cell culture and animal experiments, Emetine dihydrochloride (324693) and Gambogic acid (345701) were procured from Merck, and Celastrol (C0869) and Niclosamide (N3510) from Sigma Aldrich.

### Animal experiments

The *in vivo* anti-CHIKV activity of the test compounds was studied in the C57BL/6 mouse model of CHIKV infection (15). For the experiments, the mice (10–12 weeks old, 20–25 g, either sex) were divided into three groups, each containing 10 animals. Group 1 (mock-infected) received sterile PBS, Group 2 (CHIKV-infected) received no treatment, and Group 3 (CHIKV-infected) was administered the test compound formulated in an appropriate vehicle. ED was dissolved in sterile PBS. Celastrol and Gambogic acid were prepared in 10% DMSO diluted with sterile PBS, and Niclosamide was dissolved in 0.5% Carboxymethyl cellulose (CMC) in sterile PBS. For the virus infection, Group 2 and Group 3 mice were subcutaneously injected with a non-lethal dose of CHIKV (IND-06-Guj strain, 10^4^ pfu) in 50 µl PBS on the ventral side of both hind limbs. The CHIKV infection led to detectable blood viremia between 2–4 days post-infection (pi), peaking at 2 or 3 days pi, and paw edema starting at 2–3 days pi, peaking at 6–8 days pi. The test compounds were administered intraperitoneal or subcutaneously at different times pi. The mice were monitored for two weeks with daily measurements of body weight, and paw edema using a Vernier Calliper (paw width in mm) or digital plethysmometer (paw volume in cc). The blood collected from the animals on 2 and 3 days pi was used to determine the CHIKV RNA levels, while serum was isolated to quantify the viral titers.

### Virus binding and uptake assays

For the virus binding assay, ERMS cells were incubated with culture medium containing varying concentrations of ED for 1 h at 37 °C. Following the removal of the culture medium, the cells were incubated with CHIKV (MOI 1) in an ED-free medium at 4 °C for 1 h to allow the viral adsorption. The cells were washed twice with PBS, lysed using RNAiso Plus (Takara, 9109) for RNA isolation, and the viral RNA levels were quantified by qRT-PCR.

For the uptake assay, ERMS cells were incubated with culture medium containing varying concentrations of ED at 37 °C for 1 h. Following the removal of the culture medium, the cells were then incubated with CHIKV (MOI 1) in an ED-free medium for 1 h at 4 °C, followed by washing with PBS. Afterwards, viral uptake was allowed to proceed by incubating the cells at 37 °C for 1 h in DMEM with 2% FBS. The cells were then treated with trypsin (2.5 g/L) for 1 min (15) and washed with PBS to remove any extracellular virus. The cells were washed with PBS, lysed using RNAiso Plus (Takara, 9109) for RNA isolation, and the viral RNA levels were quantified by qRT-PCR.

### Quantitative Real-time PCR for the CHIKV RNA

To quantify the CHIKV RNA levels, the total RNA was extracted from the cells using RNAiso Plus (Takara, 9109), and cDNA was prepared using 500 ng RNA, random hexamers, and ImProm-II reverse transcription system (Promega, A3800). The relative abundance of viral RNA level was determined by quantitative real-time PCR (qRT-PCR) using a 2× SYBR-green reagent (Takara; RR420A) in the QuantStudio 6 Flex RT-PCR machine. The assay used a LightCycler® 96 Real-Time PCR System (Roche Life Science). The *Gapdh* levels were used as the internal housekeeping control. The PCR conditions were as follows: 94 °C for 2 min (1 cycle), 94 °C for 15 sec, 55 °C for 30 sec, 72 °C for 1 min (40 cycles). The viral RNA levels were normalised to Gapdh and calculated by the ΔΔCt (threshold cycle) method. All experiments had biological duplicates and were performed independently three or more times. The fold-change in the RNA level is represented as the mean ± SD of three or more independent experiments. The primers (5’-3’) used in the study were as follows: GAPDH: F-TGCACCACCAACTGCTTAGC; R-GGCATGGACTGTGGTCATGAG; CHIKV: F-GGCAGTGGTCCCAGATAATTCAAG; R-GCTGTCTAGATCCACCCCATACATG; SINV: F-AAAGGATACTTTCTCCTCGC; R-TGGGCAACAGGGACCATGCA; RRV: F-GCGACGGTGGATGTCAAGGAG; R-AGCCAGCCCACCTAACCCACTG.

### Strand-specific quantitation of CHIKV RNA

The positive- and negative-sense RNA of CHIKV was quantified using a strand-specific qRT-PCR assay as described previously (15,36). The total RNA isolated from the CHIKV-infected cells was employed for the cDNA synthesis by reverse transcription. Tagged (non-viral sequence) primers PtagCHIKp and NtagCHIKn were used for the cDNA synthesis from the positive- and negative-sense CHIKV RNA, respectively. As reported before, the real-time qPCR was performed using a combination of primers that bind to the non-viral tag sequence and viral strand (15). The strand-specific RNA copy numbers were determined using the positive- and negative-sense RNA-specific standard curves. To make the standard curve, the total RNA isolated from the CHIKV-infected cells was used for the cDNA synthesis using random hexamers. The PCR was done with the T7-tagged primers (15). The CHIKV nsP2 RNA was generated by *in vitro* transcription of the above PCR products. The RNA was quantified using a spectrophotometer and subjected to qPCR. The standard curve was plotted using the Ct values obtained from the range of known RNA concentrations and the calculated copy number of strand-specific CHIKV RNA.

### Western blotting

For Western blotting, cell lysates were prepared by resuspending the cells in the pre-chilled RIPA lysis buffer (150 mM NaCl, 1 mM EDTA, 50 mM Tris, 1% Triton X-100, 1 mM PMSF) containing protease inhibitors. The lysates were incubated on ice for 2 h and then centrifuged at 12,000 rpm for 15 min. The supernatant was collected, and the protein concentration was determined using the Bradford method. Electrophoresis resolved the protein sample (25 μg) on a 10% Sodium dodecyl sulphate (SDS)-polyacrylamide gel. The gel was electroblotted onto a Polyvinylidene fluoride (PVDF) membrane (MDI, India). Further, the membranes were blocked in 5% Bovine serum albumin (BSA) prepared in Tris-buffered saline with 0.1% Tween-20. This was followed by incubation with the primary antibody overnight at 4 °C. The membranes were washed with PBS having 0.1% Tween 20 (PBST) and then incubated in HRP-conjugated secondary antibody for 1 h. The membranes were again washed with PBST, and the blots were developed using enhanced chemiluminescence (ECL) detection reagents (Cytiva, RPN2209) and visualised using the ChemiDoc MP imaging system (Bio-Rad Labs Pvt. Ltd., India).

### Host cell protein synthesis

ERMS cells were pre-treated with varying concentrations of ED at 37 °C for 1 h, followed by the addition of 0.01 mg/mL puromycin as previously described (37). Cells treated with DMSO served as the vehicle control. The cells were washed 12 min later with the pre-warmed PBS (37 °C) and allowed to recover in drug-free culture medium for 30 min. The cell lysates were made and subjected to western blotting using anti-puromycin monoclonal antibody (Abcam, EPR27218-173). The blots were density scanned to quantify the puromycilated proteins using ImageJ software.

### Computational analysis

The crystal structure of CHIKV nsP2 helicase was retrieved from the RCSB Protein Data Bank (PDB ID: 6JIM, resolution 2.0 Å). The three-dimensional structure of Emetine dihydrochloride (ED) was obtained from the PubChem database (CID: 10219). Protein preprocessing was performed using Schrödinger’s Protein Preparation Wizard, which involved the addition of missing hydrogen atoms, assignment of bond orders, optimisation of hydrogen-bonding networks, and energy minimisation using the OPLS2005 force field. The ligand ED was prepared using the LigPrep module to generate its protonated, low-energy conformations, followed by desalting and stereoisomer generation where applicable. The SiteMap analysis was performed using Schrödinger to identify the potential ligand binding site(s) on the protein’s surface. A receptor grid was generated around the predicted active site of the helicase, focusing on conserved motifs essential for the ATPase activity. The molecular docking was carried out to evaluate the binding of ED to CHIKV nsP2 using Schrödinger’s Glide module in extra-precision (XP) mode (15). Flexible docking of the ligand was performed to explore potential binding orientations. The optimal binding pose was selected based on the GlideScore, interaction analysis, and visual inspection to identify the key interacting residues. Binding affinity (expressed in kcal/mol) was estimated using the docking score. Post-docking binding free energy (ΔG_bind_) was calculated using the Prime Molecular Mechanics with Generalised Born Surface Area (MMGBSA), incorporating solvation and entropy contributions via single-point energy calculations based on the Generalised Born Surface Area (GBSA) model. To evaluate the dynamic stability of ED at the potential binding site on CHIKV nsP2, molecular dynamics (MD) simulations were conducted using the Desmond module for 100 ns following standard relaxation protocols. The trajectory analysis was performed to assess interaction stability throughout the simulation, which included monitoring the root mean square deviation (RMSD) of the protein-ligand complex. The key binding residues were identified using the Ligand Interaction fingerprinting and hydrogen bond analysis tools.

### Expression and purification of CHIKV nsP2 protein

The full-length CHIKV nsP2 protein and its helicase domain were expressed and purified as described previously (15). Briefly, a modified pET14b expression plasmid encoding the gene of interest and an N-terminal His_6_-SUMO tag was used to transform *Escherichia coli* Rosetta (DE3) cells. Cultures were grown in LB to an OD_600_ of 0.6–0.8 and induced with 0.5 mM Isopropyl β-d-1-thiogalactopyranoside (IPTG) at 18 °C for 16 h. Cells were harvested and lysed in buffer containing 20 mM HEPES pH 7.5, 500 mM NaCl, 5% Glycerol and 2 mM β-Mercaptoethanol. The lysate was clarified and applied to a Ni-NTA column (Cytiva), followed by elution using an Imidazole gradient. His_6_-SUMO tag was removed using PreScission protease, and the sample was dialysed overnight at 4 °C. The protein was concentrated and subjected to size-exclusion chromatography using the Superdex 200 16/600 (Cytiva) column in 20 mM HEPES pH 7.5, 300 mM NaCl, 5% Glycerol, and 0.5 mM DTT. The purity was confirmed by SDS PAGE.

### Isothermal Titration Calorimetry (ITC)

The ED binding interactions with CHIKV nsP2 full-length (FL) protein and its truncated domains were studied by ITC using the MicroCal ITC-200 machine (MicroCal, USA). The proteins and ED were prepared in a buffer containing 20 mM HEPES (pH 7.5), 250 mM NaCl, 5% Glycerol, and 1% DMSO. The calorimetric cell was loaded with 10 µM protein, while 100 µM ED was loaded into the syringe for titration into the reaction cell. The titration consisted of an initial 0.4 µl injection, followed by 18 injections of 2 µl at 150 sec intervals, with an initial delay of 60 sec. The titration was conducted at 25 °C with a reference power of 10 µcal/s and a stirring speed of 750 rpm. The raw ITC data and titration plots were analysed using Malvern’s Origin 7.0 Microcal-ITC200 analysis software, employing a one-site binding model to calculate the thermodynamic parameters, including the dissociation constant (K_d_), and enthalpy change (ΔH). The thermodynamic changes due to ED dilution were negligible when the buffer was titrated against ED and vice versa. The binding interactions were characterised by significant changes in differential power (DP) in µcal/sec and binding enthalpy (ΔH) profiles, as shown in the isotherms, with the solid lines representing the best least-squares fitting to the data.

### Microscale Thermophoresis (MST)

The ED binding with the CHIKV nsP2 protein was studied by MST on the Monolith NT.115 instrument (NanoTemper) as per the manufacturer’s instructions. For the MST experiments, 10 µM protein was labelled on Lysine residues with RED-NHS fluorescent dye using the 2nd Generation Monolith Protein Labelling RED-NHS kit (NanoTemper). The dye contains a reactive NHS-ester group that forms a covalent bond with the primary amines, such as Lysine residues, by reacting with them. For binding affinity, the labelled protein (1 µM) was incubated with two-fold serial dilutions of ED (starting from 250 µM) and loaded into the Monolith NT.115 capillaries to obtain the binding curve. The data were collected using the Monolith NT.115 device, with 60% (RED) LED power, and analysed using MO Affinity Analysis v2.3 software. For binding affinity analysis, ligand-dependent changes in temperature-related intensity change (TRIC) were plotted as Fnorm values against ligand concentration in a dose-response curve. The Fnorm values were expressed in parts per thousand (‰). For each trace, the Fnorm value was determined by dividing F1 by F0, where F1 represents the fluorescence value measured in the heated state, and F0 corresponds to the fluorescence value measured in the cold state before the IR laser is activated. The MST data showed an affinity curve with well-defined bound and unbound states. The dissociation constant (K_d_) for the interaction of CHIKV nsP2 and its truncated helicase domain with ED was estimated.

### Site-directed mutagenesis

The polymerase chain reaction (PCR) was used for the site-directed mutagenesis employing overlapping primers on the expression plasmid containing the CHIKV nsP2 cDNA. The overlapping primers were designed according to the desired nucleotide changes in the coding region of the protein. The PCR was carried out using Phusion polymerase (NEB, M0530S) and the reaction mix was digested with DpnI (NEB, R0176S) to digest the methylated parent plasmid (the template). The resultant product DNA was purified and used to transform the E. coli DH5α competent cells. The cells were grown at 37 °C, and the plasmid was isolated. It was subjected to nucleotide sequencing to verify the cDNA sequence with the desired mutation.

### Helicase assay

To investigate the effect of ED on RNA unwinding activity of CHIKV nsP2, a helicase assay was performed following the protocol described previously (38,39). The assay utilised an Alexa Fluor 488-labelled 28-mer ssRNA oligonucleotide (Alexa488-ssRNA, 5’-AAAAAAAAAAAACCAGGCGACAUCAGCG-3’) and an unlabelled 16-mer ssRNA oligonucleotide (5’-CGCUGAUGUCGCCUGG-3’) to generate a dsRNA helicase substrate with a 12-base 5’ overhang. To prepare the dsRNA substrate, the labelled and unlabelled RNA oligonucleotides were mixed at a 1:1.1 molar ratio in a buffer containing 10 mM HEPES (pH 7.2) and 20 mM KCl. To facilitate annealing, the RNA mixture was heated to 95 °C for 1 min in a thermal cycler, followed by gradual cooling at a rate of 1 °C per min until reaching 22 °C. The optimized reaction mixture for the unwinding assay consisted of the strand-displacement assay buffer (40 mM HEPES, pH 7.5, 2 mM dithiothreitol (DTT), and 12 mM NaCl), 1 μM purified CHIKV nsP2 protein or its helicase and protease domains, 50 nM dsRNA substrate, 800 nM unlabelled RNA trap (5’-CCAGGCGACAUCAGCG-3’), 20U RNaseOUT inhibitor (Thermo Fisher Scientific, 10777019), and varying concentrations of ED. The reaction mixture was incubated at 26 °C for 20 min in a thermal cycler to facilitate the formation of the protein-RNA complex. Subsequently, a 3.5 mM ATP-magnesium acetate mixture was added, followed by incubation at 37 °C for 120 min. The reactions were then terminated by adding a stop solution containing 100 mM Tris-HCl, pH 7.5, 0.1% bromophenol blue, 1% SDS, 50 mM EDTA, and 50% Glycerol. The separation of ssRNA and dsRNA was carried out through electrophoresis on a 15% nondenaturing polyacrylamide gel at 4 °C. The RNA bands were visualized using a Typhoon imager (Cytiva), and their intensities were quantified using the ImageJ software. Denatured ssRNA (prepared by heating dsRNA at 95 °C for 5 min, followed by rapid cooling at 4 °C) was used as a control for dsRNA. BSA protein (1 μM) was used as a negative control to assess the unwinding activity of nsP2.

## RESULTS

### Screening of the Spectrum Collection for the anti-CHIKV compounds

BHK-21 cells were used for the screening assay, for which various conditions, such as the cell density, viral-infective dose as the multiplicity of infection (MOI) and assay endpoint, were standardised. Ribavirin was used as the positive control since it has been shown to inhibit CHIKV *in vitro* and *in vivo* (15). To ensure a reliable conclusion, validation of the assay was done by studying the quality parameters; the signal-to-noise (S/N) ratio was ∼50, the coefficient of variation (CV) was <1.9%, and the Z’ factor was 0.91.

In the primary screening of the Spectrum Collection in BHK cells, 164 compounds showed >80% cell viability and >80% CHIKV inhibition at 10 µM concentration (Fig. 1A). These compounds were screened in the second round at 1 µM concentration in ERMS cells where 4 compounds showed >80% cell viability and >80% CHIKV inhibition (Fig. 1A). These were Celastrol, Niclosamide, Gambogic acid, and Emetine dihydrochloride (ED). The CC_50_ and IC_50_ of these compounds were determined in ERMS cells to calculate the selectivity index (SI) (Fig. 1B). The SI value for Celastrol was 123, for Niclosamide it was 65, and for Gambogic acid it was 2. ED had a very high SI value of 967.

**Figure 1:**
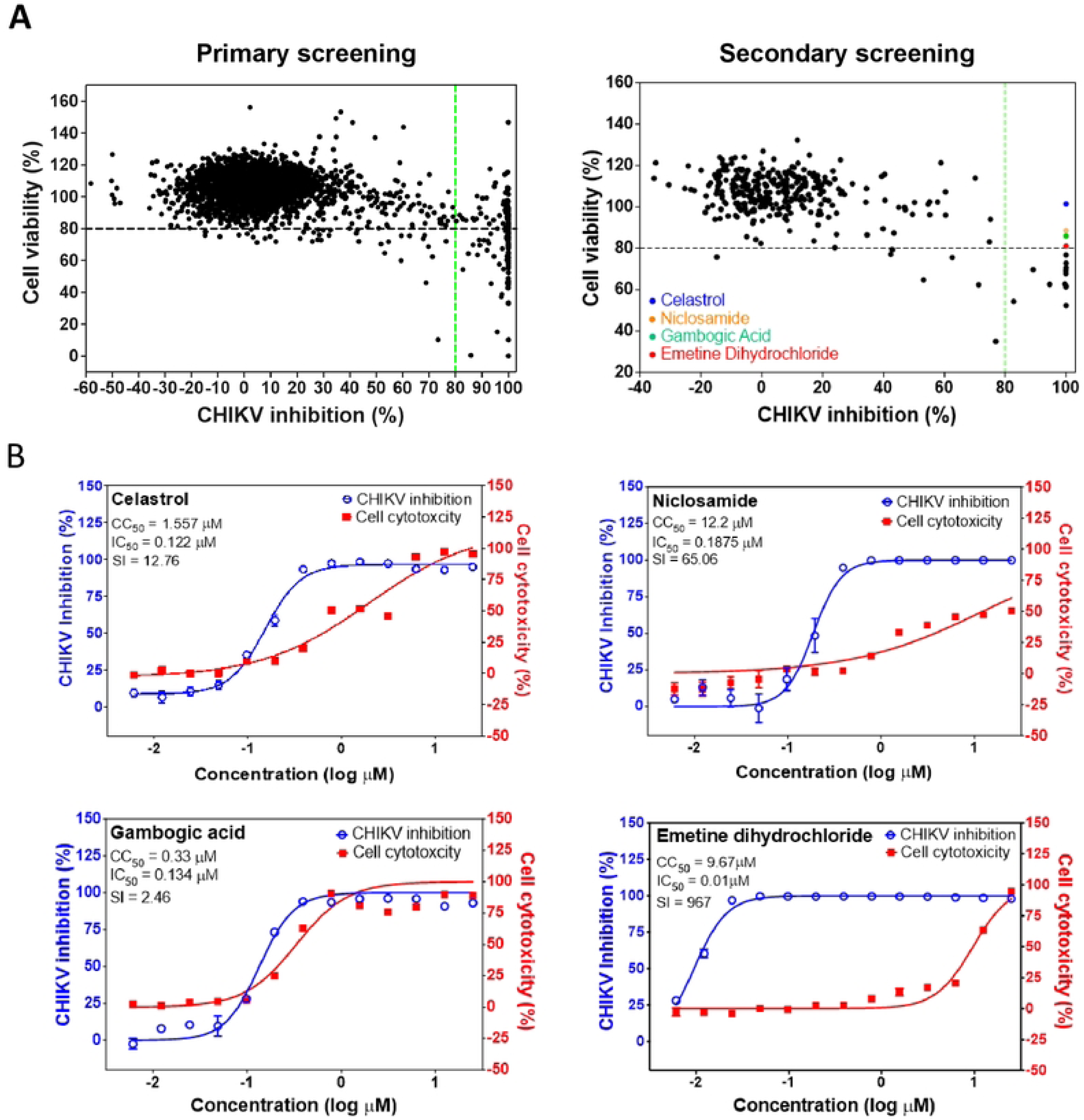
Screening of the Spectrum Collection for the CHIKV antiviral activity. (A) BHK-21 (for primary screening) or ERMS (for secondary screening) cells were seeded in a 96-well plate and infected with CHIKV-GFP at 0.1 or 5 MOI for 20 or 32 h, respectively, and treated with DMSO (vehicle control) or the test compounds. The left panel (primary screening) shows the cell viability and CHIKV replication inhibition in BHK cells using different compounds at a 10 µM concentration. The right panel (secondary screening) shows the cell viability and CHIKV replication inhibition in ERMS cells using different compounds at 1 µM concentration. (B) ERMS cells were infected with CHIKV-GFP (5 MOI) and treated with different concentrations of the test compounds. The dose-response curve demonstrating the cell toxicity and CHIKV inhibition at different concentrations at 24 h pi is shown.

### Anti-CHIKV efficacy of the selected compounds in mice

The antiviral potential of Celastrol, Niclosamide, Gambogic acid, and ED was tested in the mouse model of CHIKV infection in adult C57BL/6 mice, where CHIKV-infected mice show viremia and footpad edema, which is self-resolving (Fig. 2). Mice inoculated subcutaneously in the footpad with CHIKV began to show footpad edema at 4-5 day pi, with maximum edema recorded at 6-7 day pi, after which the edema was seen to self-resolve reaching the normal levels by 10-12 day pi. CHIKV-infected mice were treated with the above compounds at a dose of 10 mg/kg intraperitoneal, 4 h after the virus infection. Treatment of CHIKV-infected mice with Niclosamide or Gambogic acid did not affect the virus-induced footpad edema. Celastrol treatment delayed the footpad edema without affecting its extent. ED-treated CHIKV-infected mice, however, showed no edema (*p*<0.001). The footpad edema measurements of the ED-treated CHIKV-infected mice were statistically not different from those in the mock-infected mice.

**Figure 2.**
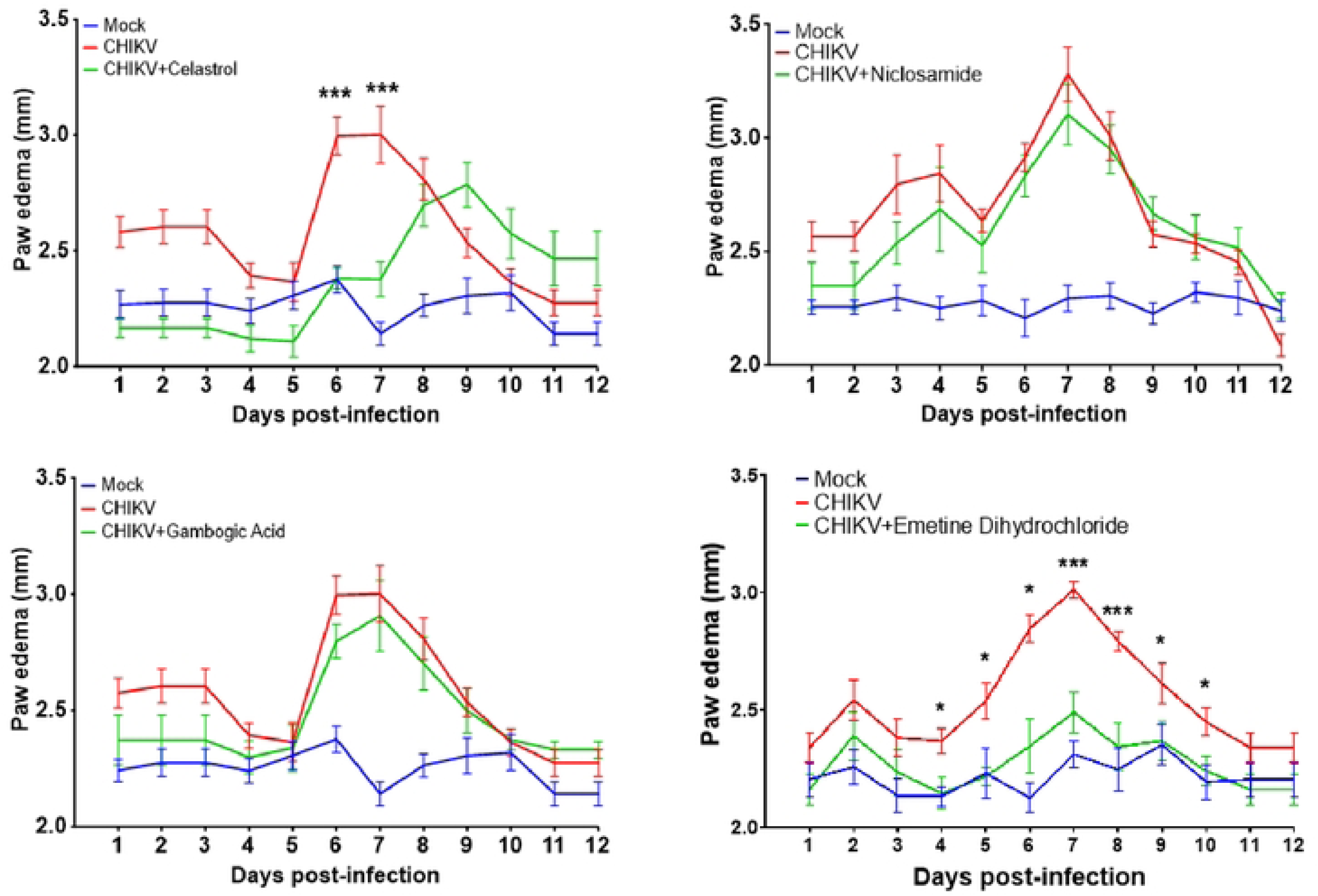
Anti-CHIKV efficacy of the selected compounds in the mouse model of disease. C57BL/6 mice of 12 weeks of age were mock-infected or infected subcutaneously with 10^4^ pfu of CHIKV and treated with vehicle alone or the indicated drug compound (5 mg/kg) dissolved in an appropriate solvent, given intraperitoneal once a day. The first dose of the compound was delivered 4 h pi. The mice were followed for 12 days, and the paw edema was measured daily as paw width/thickness using a Vernier Calliper. A line graph demonstrating the mouse paw edema on different days pi is presented. The Bonferroni post-hoc test, followed by a two-sided independent t-test, was used to calculate the *p* values: \**p*<0.0332, \*\**p*<0.0021, \*\*\**p*<0.0002, \*\*\*\**p*<0.0001.

We then studied the efficacy of ED, delivered subcutaneously at lower doses, in CHIKV-infected mice (Fig. 3A). A dose effect was seen on the virus-induced footpad edema in CHIKV-infected mice treated with graded doses of ED. CHIKV-infected mice treated with the 6 mg/kg dose of ED showed no footpad edema. Mice treated with the 3 mg/kg dose of ED showed significantly reduced footpad edema compared to the control. However, ED at the 1 mg/kg dose did not reduce the virus-induced footpad edema in CHIKV-infected mice. Similarly, there was no weight loss in the CHIKV-infected mice treated with a 6 mg/kg dose of ED, whereas weight loss wasn’t fully recovered in CHIKV-infected mice treated with ED at a dose of 3 mg/kg or 1 mg/kg (Fig. 3B). The antiviral effect of ED in CHIKV-infected mice was reflected in the significantly reduced CHIKV viremia (Fig. 3C). Here also, a dose-response was seen in the reduced CHIKV titer in mice treated with the graded doses of ED, with the highest reduction recorded in mice treated with the 6 mg/kg dose.

**Figure 3.**
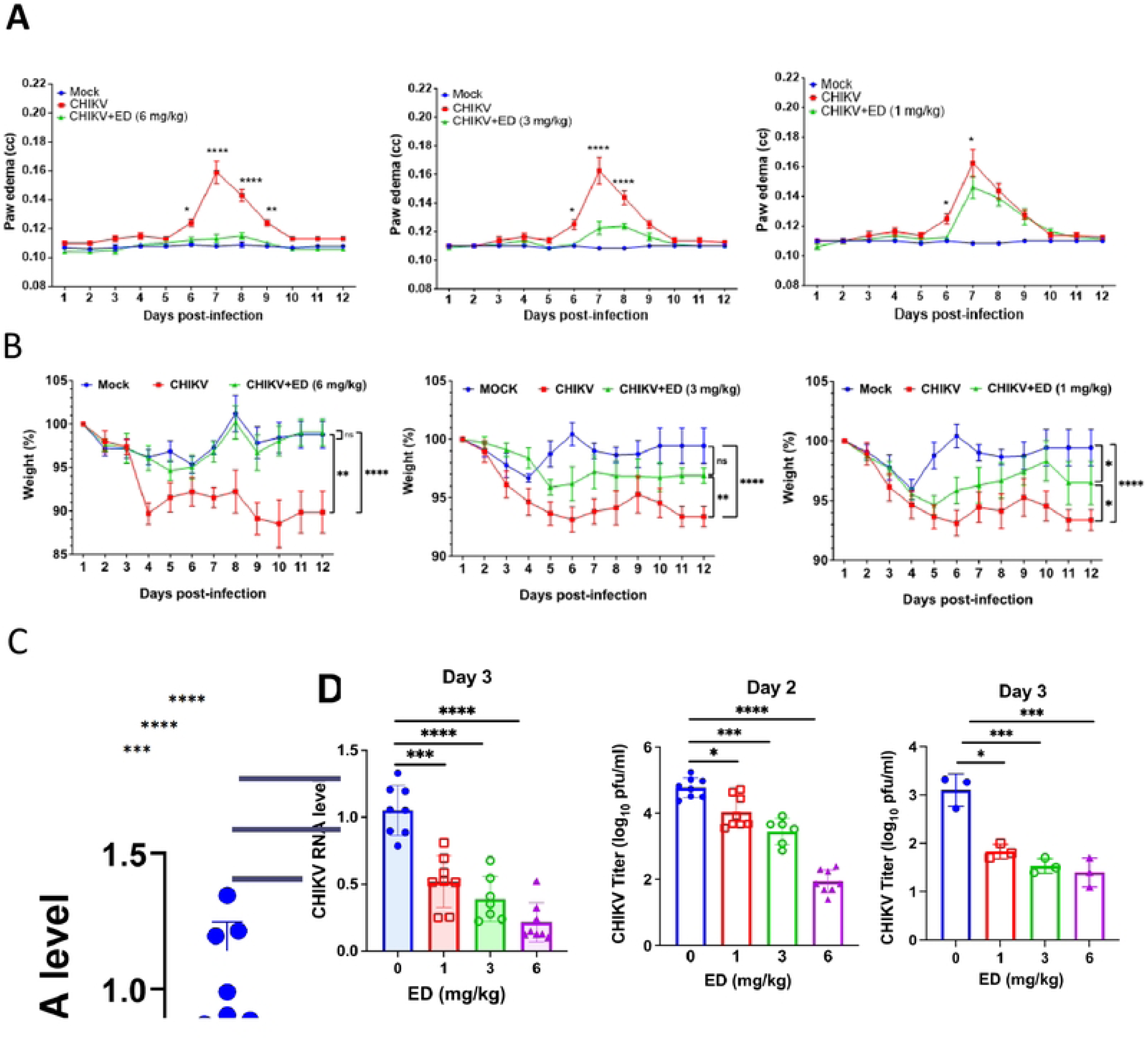
Anti-CHIKV efficacy of ED in the mouse model of disease. C57BL/6 mice of 12 weeks of age were mock-infected or infected subcutaneously with 10^4^ pfu of CHIKV and treated with vehicle alone or the indicated dose of ED given subcutaneously once a day. The first dose of the compound was delivered 4 h pi. The mice were followed for 12 days, and the paw edema was measured daily using a digital plethysmometer. (A) A line graph demonstrating the mouse paw edema on different days pi is presented. The Bonferroni post-hoc test, followed by a two-sided independent t-test, was used to calculate the *p* values: \**p*<0.0332, \*\**p*<0.0021, \*\*\**p*<0.0002, \*\*\*\**p*<0.0001. (B) A line graph demonstrating the mouse weight on different days pi is presented. The Bonferroni post-hoc test, followed by a two-sided independent t-test, was used to calculate the *p* values: \**p*<0.0332, \*\**p*<0.0021, \*\*\**p*<0.0002, \*\*\*\**p*<0.0001. (C) Blood was drawn from the animals on 2 and 3 days pi and used to determine the CHIKV titers and RNA levels. The relative CHIKV RNA levels (compared to the no ED-treatment control) and viral titers are presented. The student’s t-test was used to calculate the *p* values: \**p*<0.05, \*\**p*<0.01, \*\*\**p*<0.001, \*\*\*\**p*<0.0001.

### Antiviral potential of ED against CHIKV in mice

To test the antiviral potential of ED to treat CHIKV patients, CHIKV-infected mice were treated with ED at different times pi. ED treatment was effective in suppressing the virus-induced footpad edema when delivered at 4 h pi, 24 h pi, and as late as 48 h pi (Fig. 4A). A similar effect of ED was seen on the mouse weight (Fig. 4A). Again, ED treatment led to a reduced viremia in CHIKV-infected animals (Fig. 4B). Drug treatment beyond 48 h pi was not investigated.

**Figure 4.**
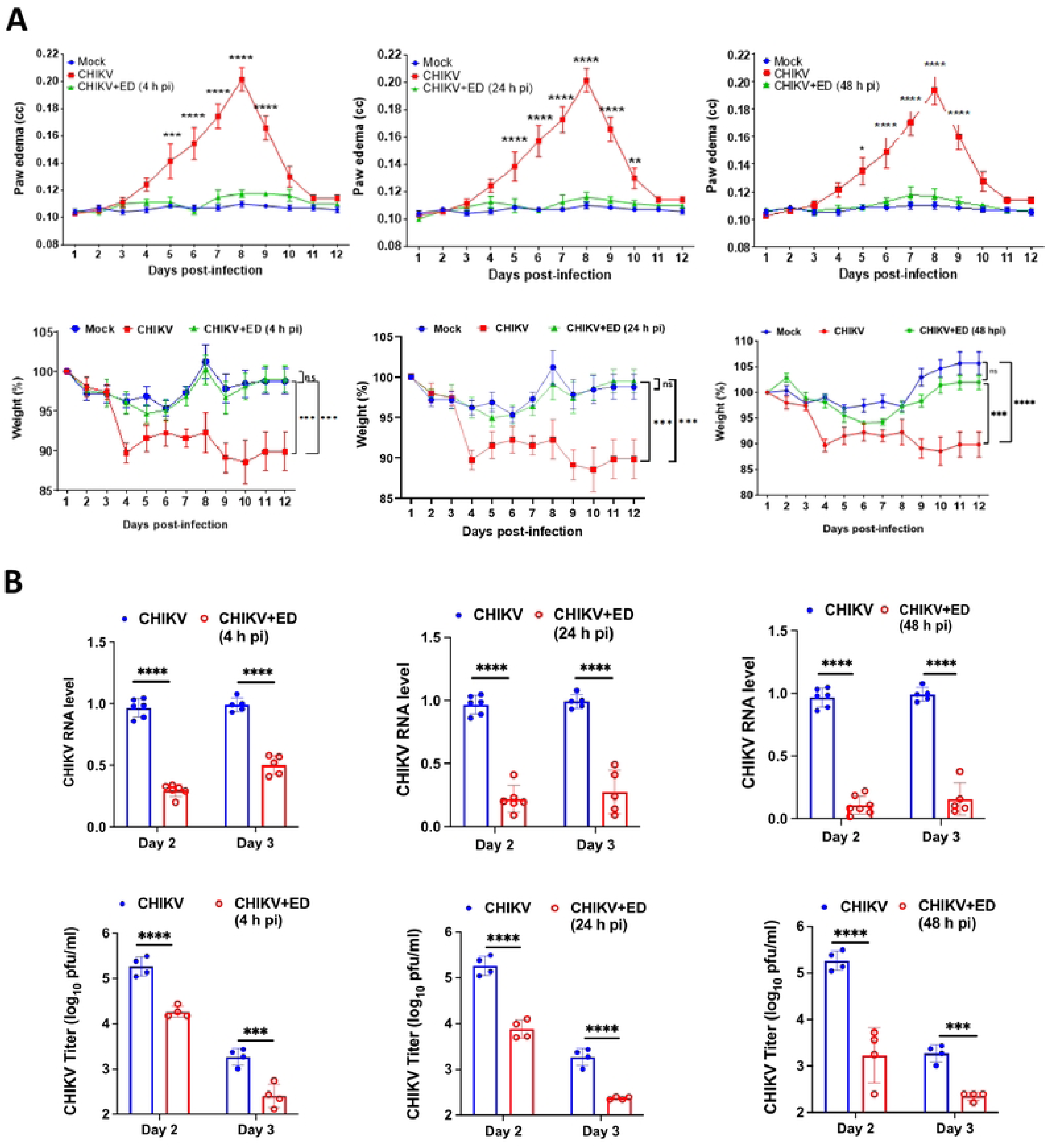
Antiviral potential of ED in the mouse model of Chikungunya disease. C57BL/6 mice of 12 weeks of age were mock-infected or infected subcutaneously with 10^4^ pfu of CHIKV and treated with vehicle alone or ED (6 mg/kg) given subcutaneously once a day. The first dose of the compound was delivered at 4, 24, or 48 h pi. The mice were followed for 12 days, and the paw edema was measured daily using a digital plethysmometer. (A) Line graphs in the top panel demonstrating the mouse paw edema on different days pi are presented. The bottom panel has the line graphs demonstrating the mouse weight on different days pi. The Bonferroni post hoc test, followed by a two-sided independent t-test, was used to calculate the *p* values: \**p*<0.0332, \*\**p*<0.0021, \*\*\**p*<0.0002, \*\*\*\**p*<0.0001. (B) Blood was drawn from the animals on 2 and 3 days pi and used to determine the CHIKV titers and RNA levels. The relative CHIKV RNA levels (compared to the no ED-treatment control) and viral titers are presented. The student’s t-test was used to calculate the *p* values: \**p*<0.05, \*\**p*<0.01, \*\*\**p*<0.001, \*\*\*\**p*<0.0001.

### CHIKV replication kinetics in ERMS cells in the presence of ED

To understand the mechanism of ED’s antiviral action, the CHIKV replication kinetics was studied in the presence of various ED concentrations. The CHIKV genome replication, as determined by the CHIKV RNA levels, was significantly suppressed in the ED-treated CHIKV-infected cells as early as 3 h pi and this suppression continued till 12 h pi (Fig. 5). The viral RNA levels were lower in cells treated with ED concentration as low as 0.01 µM. The viral titers increased with time in CHIKV-infected control cells. However, these were significantly lower in the presence of ED concentration as low as 0.01 µM and 0.1 µM at 6 and 12 h pi, respectively. Importantly, the viral RNA replication and the viral titers in the presence of varying ED concentrations showed a dose effect.

**Figure 5.**
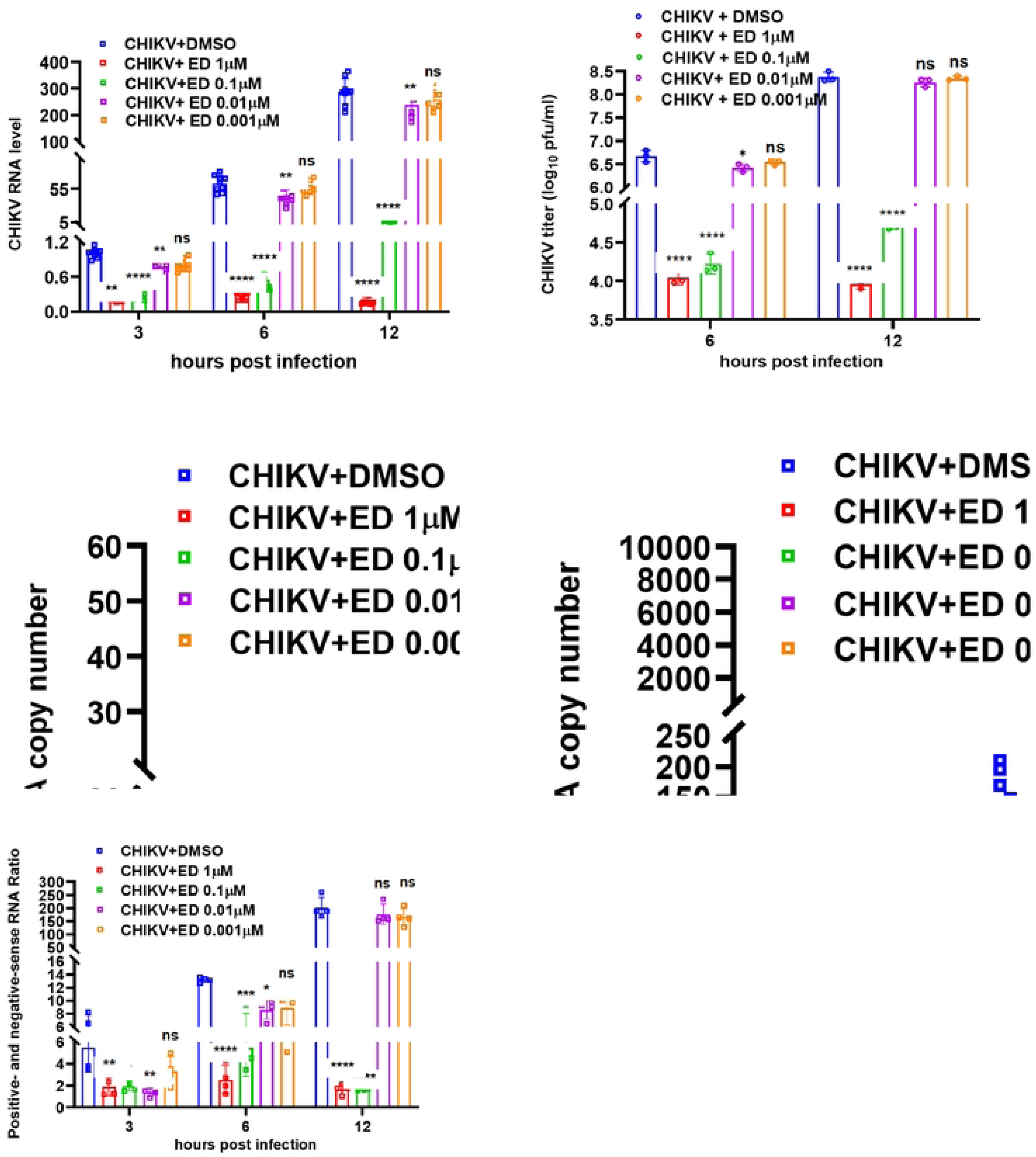
CHIKV replication kinetics in ERMS cells in the presence of ED. ERMS cells were infected with CHIKV (1 MOI) and incubated with different concentrations of ED. The control CHIKV-infected cells were incubated with DMSO. The cells and culture supernatants were harvested at different times pi for the extraction of the total RNA and determination of the viral titers, respectively. The qRT-PCR was used to determine the CHIKV RNA levels and the positive- and negative-sense CHIKV genome copy numbers. The relative viral RNA levels are shown in the top left panel, where *Gapdh* was used as the internal control. The RNA level in the control at 3 h pi was taken as one. The CHIKV titers determined by plaque assay are shown in the top right panel. The negative- and positive-sense CHIKV genomic RNA copy numbers are shown in the middle panel. The bottom panel shows the ratio of the positive- and negative-sense RNAs. The data in the control (DMSO-treated) cells were compared with those treated with ED at different time points. The student’s t-test was used to calculate the *p* values: \**p*<0.05, \*\**p*<0.01, \*\*\**p*<0.001, \*\*\*\**p*<0.001.

In the CHIKV-infected control cells, the negative-sense CHIKV RNA could be detected at 3 h pi, and its copy number continued to increase till 12 h pi (Fig. 5). In the CHIKV-infected ED-treated cells, however, the negative-sense CHIKV RNA levels were significantly lower in the presence of ED concentration as low as 0.01 µM (Fig. 5). Similarly, the positive-sense RNA levels were also significantly lower in the presence of ED concentration as low as 0.01 µM. While the ratio of the positive- and negative-sense RNA showed a sharp increase in the CHIKV-infected control cells, this was not seen in the cells treated with ED concentrations as low as 0.1 µM (Fig. 5), indicating a significant inhibition of CHIKV RNA synthesis.

### ED inhibits CHIKV nsP2 protein synthesis

The reduced negative-sense RNA levels in the presence of ED early during the infection suggested that the drug inhibited the early step/s of the CHIKV life cycle. During the early phase of infection, the RNA genome of CHIKV is translated into a polyprotein, which is subsequently processed to produce the non-structural proteins required for the synthesis of the negative-sense genomic intermediate RNA that acts as a template for genome replication. The CHIKV nsP2 is a multifunctional non-structural protein with the C-terminal cysteine protease and N-terminal helicase activity-containing domains (40). It plays a crucial role in the cleavage of the polyprotein required for producing the essential viral proteins, including nsP4, with the RNA-dependent RNA polymerase activity necessary for the synthesis of the negative-sense genomic RNA and replicating the viral genome. Besides, the helicase activity is essential for the viral genomic RNA replication (38,41).

Synthesis of the CHIKV nsP2 protein was studied in virus-infected ERMS cells in the presence of the graded doses of ED (Fig. 6A). Western blotting showed inhibition of the nsP2 protein synthesis in the presence of ED at a dose as low as 0.1 µM. However, the protein synthesis inhibition was waning at further lower doses of ED.

**Figure 6.**
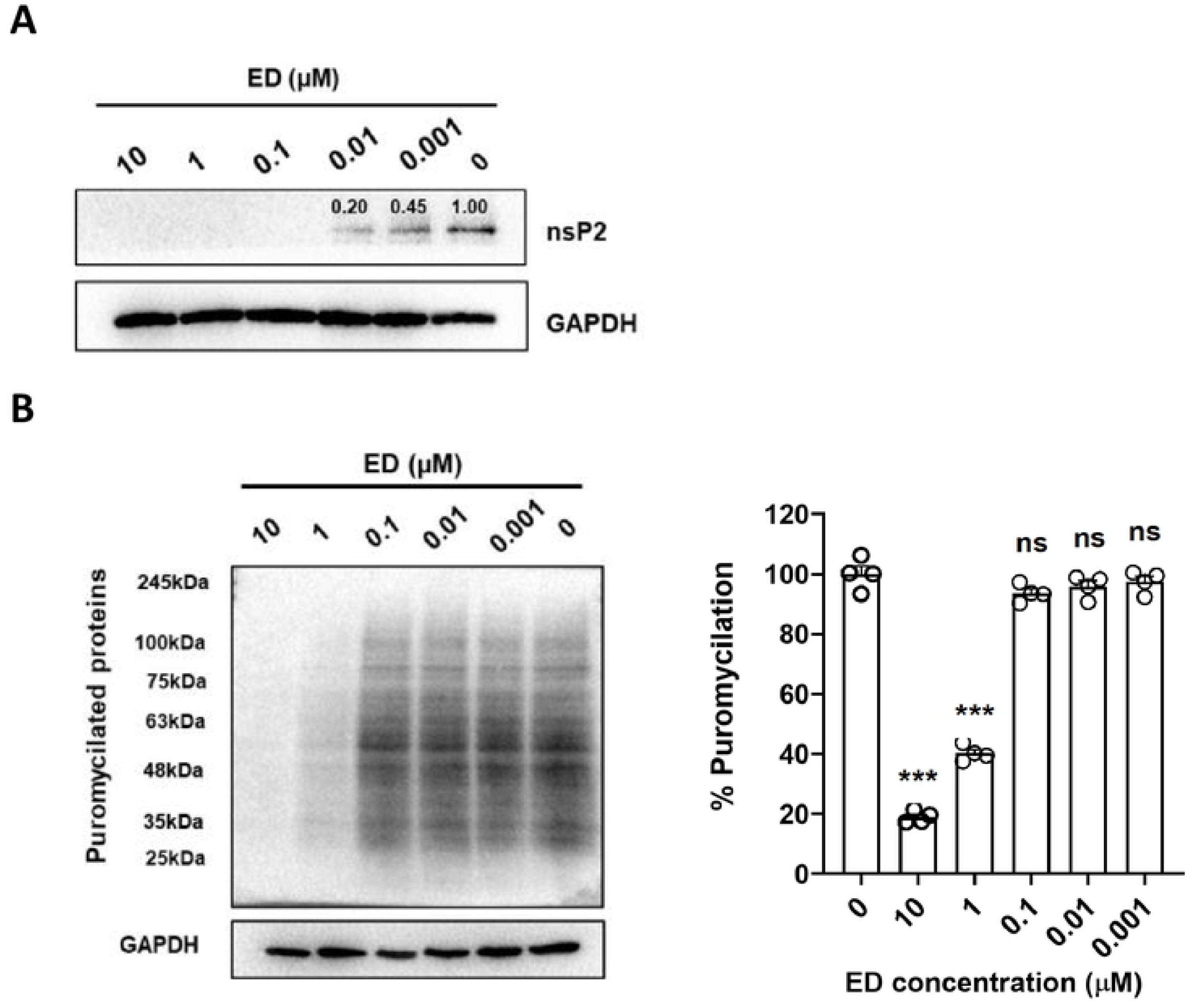
ED inhibits CHIKV protein synthesis. (A) ERMS cells were infected with CHIKV (MOI 1) and treated with different concentrations of ED, while control CHIKV-infected cells were treated with DMSO only. Cell lysates were collected at 12 h pi and western blotted with the CHIKV nsP2 antibody. GAPDH was used as the loading control. Relative band intensity was quantified using ImageJ software and displayed over the bands. (B) ERMS cells were incubated with 2% FBS-DMEM containing varying concentrations of ED for 1 h at 37 °C, followed by the addition of puromycin at a final concentration of 0.01 mg/mL. Cells treated with culture media supplemented with DMSO, without ED, served as the control. The cells were washed 12 min later with pre-warmed PBS (37 °C) and allowed to recover in 2% FBS-DMEM for 30 min. The cell lysates were western blotted with anti-puromycin monoclonal antibody. A representative Western blot is shown at the left. GAPDH was used as the loading control. The lane intensities, measured by ImageJ, relative to the control, are shown at the right. The student’s t-test was used to calculate the *p* values: \**p*<0.05, \*\**p*<0.01, \*\*\**p*<0.001, ns = not significant.

We then studied whether ED inhibited the cellular protein synthesis at concentrations where the CHIKV nsP2 protein synthesis was inhibited (Fig. 6B). We performed puromycin labelling of the host cell proteins to track the formation of puromycylated peptides in the presence or absence of ED. Puromycin integrates into the nascent polypeptide chains, prematurely terminating translation and generating puromycylated peptides. These peptides can be detected using western blotting with an anti-puromycin antibody. The host cell protein synthesis was almost completely inhibited at 10 µM ED concentration, and it was partially inhibited at 1 µM ED concentration. However, at 0.1 µM or lower ED concentration, the host cell protein synthesis remained unaffected. This suggests that the ED-mediated CHIKV protein synthesis inhibition mechanism may be different from the known ribosome-mediated inhibition of cellular protein synthesis by ED (42–44).

### ED binds to the CHIKV nsP2 protein *in silico*

Since ED inhibited the CHIKV RNA replication, and the nsP2 helicase activity is essential for the viral RNA synthesis, we examined *in silico* if ED could bind to the CHIKV nsP2 helicase domain to inhibit its activity. The SiteMap analysis predicted five potential binding sites in the helicase domain of nsP2 (Table S1). Site-1, with a site score of 1.019, was selected for molecular docking of ED (Figs. 7A, 7B). Incidentally, this site overlaps with the active site of the helicase domain of nsP2 (41). ED binds to Site-1 with a docking score of -5.399 kcal/mol. In addition, thermodynamic calculation was performed on docked poses using the prime MMGBSA program of the Schrodinger suite, which showed ED’s strong binding to Site-1 with a binding free energy (ΔG_bind_) value of -61.77 kcal/mol. The docking analysis revealed that ED was positioned in a well-defined orientation within Site-1, forming stable interactions with the key residues of the nsP2 protein helicase active site (41).

**Figure 7:**
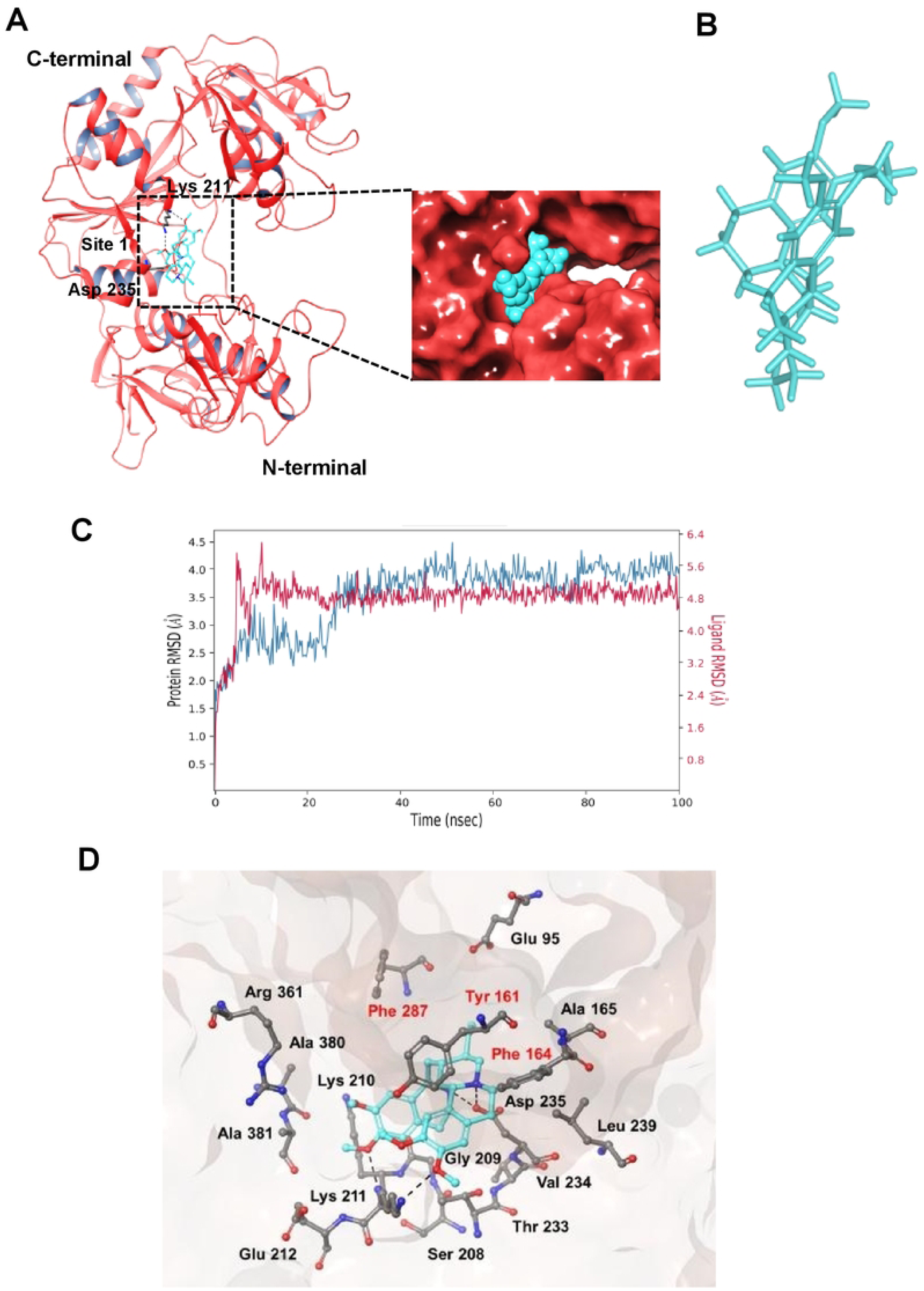
ED binding to CHIKV nsP2 helicase *in silico*. (A) The structure of the CHIKV nsP2 helicase protein (PDB: 6JIM) with docked ligand (ED) is shown. The protein is rendered in the cartoon representation and coloured in red, while ED is shown in the stick representation and coloured in cyan. The binding of ED at Site-1 is shown. The hydrogen bonding residues of nsP2 are represented in stick in grey. The inset shows the fitting of ED in the respective pocket; the protein is rendered in the surface view in red colour, and ED is shown in the vdW surface representation. (B) The structure of the ligand (ED) is shown in the stick representation. (C) The RMSD plot of the CHIKV nsP2 protein and ED interaction at Site-1 from the MD analysis is shown. (D) The interacting residues of nsP2 with ED within the Site-1 within 4.0 Å distance are shown. The interaction between the ligand and the protein is shown with black dotted lines. The protein and the ligand are in the stick representation. Amino acids involved in the substrate binding are indicated in red text.

Further, to check the conformational stability of the nsP2-helicase-ED complex, the MD simulations were carried out for 100 ns. The RMSD analysis of the MD trajectories showed that ED was stable at Site-1, and its binding stabilised the overall structure of the nsP2 protein till 100 ns (Fig. 7C). The MD simulation outcomes corroborated the docking findings that ED could bind to Site-1 on CHIKV nsP2 helicase domain.

The interaction fingerprinting identified the key amino acid residues contributing to the most stable state obtained from the MD simulation (Fig. 7D). Nine amino acids of the nsP2 helicase domain line ED within the 4.0 Å distance. The major contributors included positively charged (Lys210, Lys211), negatively charged (Asp235), hydrophobic (Phe286, Phe287, Phe164), non-polar (Ala381), and polar (Thr261, Tyr161) amino acids (Fig. 7D). Among these, Lys211 and Asp235 formed hydrogen bonds with ED, while Lys210 showed pi-pi stacking interactions with ED. These data indicate that ED has the potential to interact with the key residues (Tyr161, Phe164, and Phe287) of the substrate binding site/active site of CHIKV nsP2 helicase described previously by others (41,45).

### ED binds to the CHIKV nsP2 helicase domain

Isothermal Titration Calorimetry (ITC) was employed to study the binding of the CHIKV nsP2 protein with ED (Fig. 8A). ED was found to bind the full-length nsP2 protein in a dose-dependent manner, with a K_d_ value of 1.2 µM. We then evaluated the ED binding with the helicase and protease domains of the nsP2 protein. Here, ED was seen binding with the helicase domain in a dose-dependent manner, although the binding affinity of 2.56 µM was lower than that seen with the full-length nsP2 protein. ED showed no/very poor binding with the protease domain.

**Figure 8.**
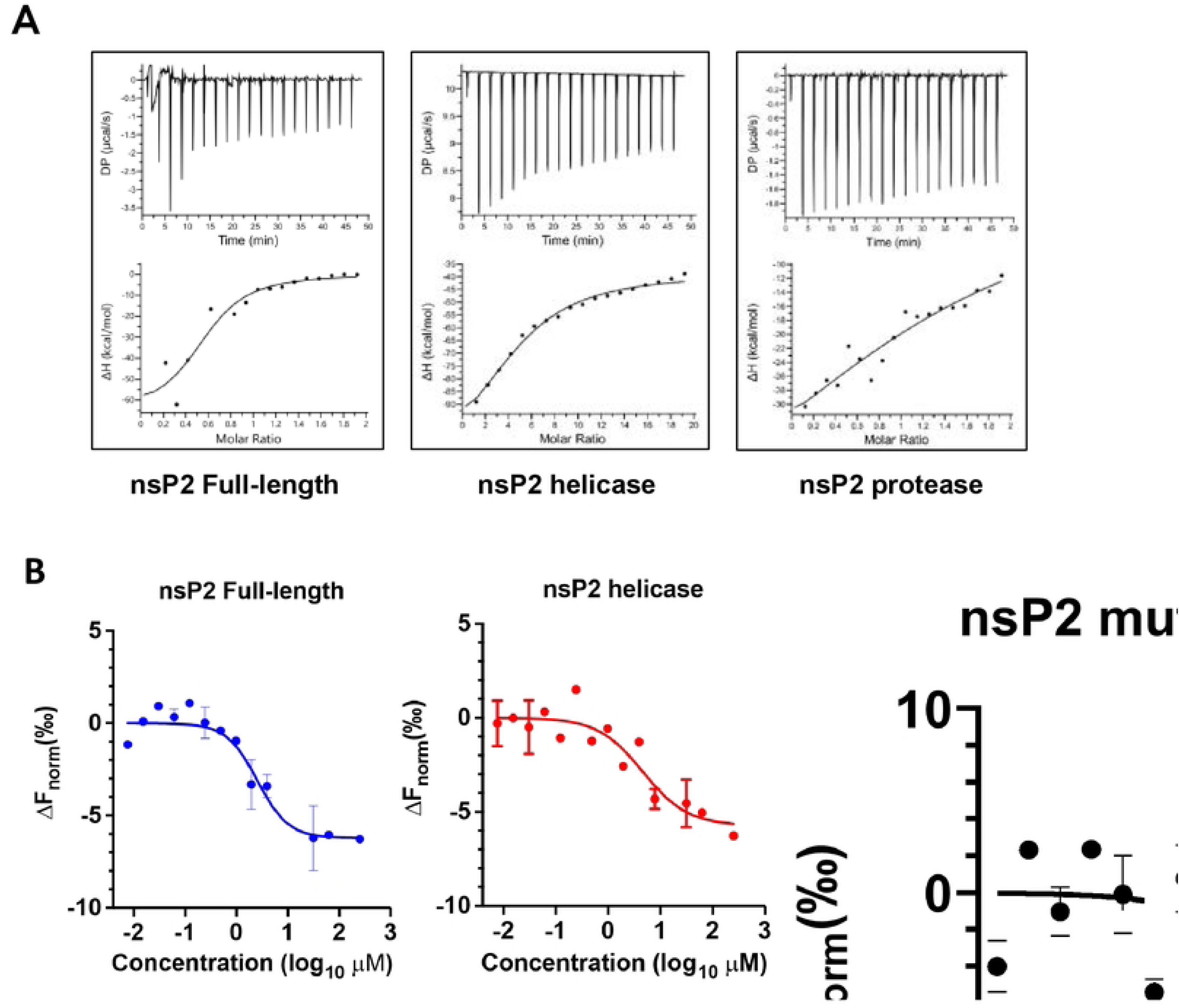
ED binds to the CHIKV nsP2 helicase domain. (A) ITC was used to study the binding of the CHIKV nsP2 protein or its helicase and protease domains with ED. The binding reactions were titrated using 10 µM protein and 100 µM ED. The thermograms (top) and fitted binding isotherms (bottom) for the binding of the nsP2 full-length protein and its truncated domains with ED are shown. Malvern’s Origin 7.0 Microcal-ITC200 analysis software was applied to obtain the dissociation constant (K_d_) and thermodynamic parameter ΔH. (B) MST was used to study the binding of the labelled CHIKV nsP2 protein (1 μM) or its helicase and protease domains with different concentrations of ED. For the binding affinity analysis, ligand-dependent changes in temperature-related intensity change (TRIC) are plotted as Fnorm values against the ED concentrations in a dose-response curve. The Fnorm values are plotted as parts per thousand (‰).

To confirm these findings further, ED binding with nsP2 was also studied by Microscale Thermophoresis (MST) (Fig. 8B). A dose-dependent binding of ED was seen with full-length nsP2 with a K_d_ value of 1.5 µM. Here also, ED bound to the helicase domain with a lower affinity, with a K_d_ of 3.06 µM. To validate the ED binding site predicted *in silico,* a Phe164Ala mutant of nsP2 was made, and its binding with ED was evaluated. The mutant showed a K_d_ value of 29.40 µM, indicating a significant reduction in binding of the mutant protein with ED compared to the wild-type nsP2.

### ED inhibits the nsP2 helicase activity

The CHIKV nsP2 helicase activity was examined using an Alexa488-tagged synthetic dsRNA. The full-length nsP2 was able to efficiently unwind the dsRNA to produce the Alexa488-tagged ssRNA (Fig. 9A). The nsP2 helicase domain also showed unwinding activity, although it was less efficient than the full-length nsP2. The nsP2 protease domain, however, did not show any helicase activity. BSA used as a negative control showed no detectable unwinding activity. The effect of ED on nsP2 helicase activity was studied using this assay. ED inhibited the helicase activity of the nsP2 protein in a dose-dependent manner (Fig. 9B). A significant inhibition of the helicase activity was seen at 0.1 µM ED concentration, at which the virus RNA and protein synthesis, and the viral replication were inhibited.

**Figure 9.**
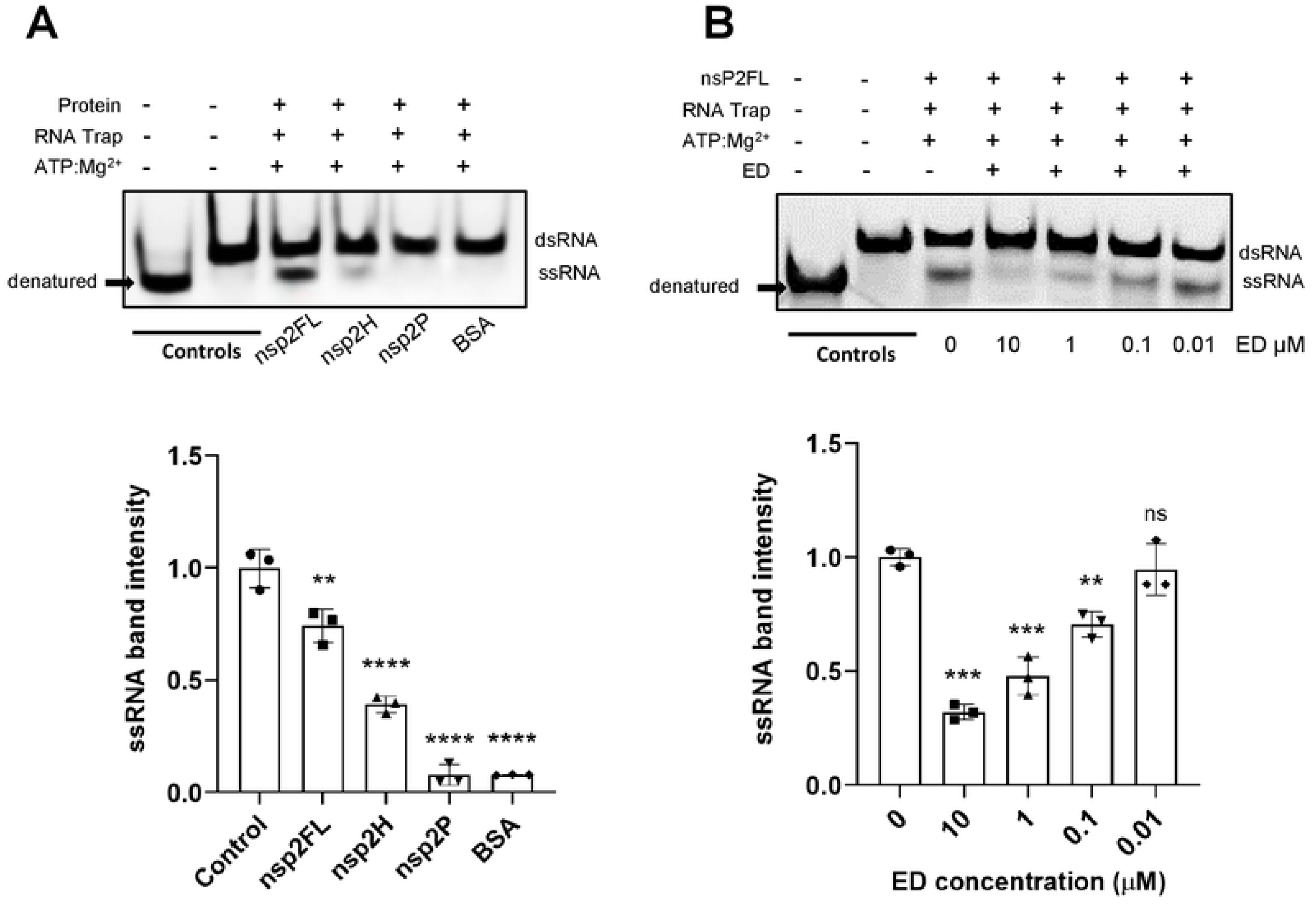
ED inhibits CHIKV nsP2 RNA Helicase Activity. (A) The RNA helicase activity of the full-length nsP2 (nsP2FL) and its helicase (nsP2H) and protease (nsP2P) domain proteins was studied using the Alexa488-dsRNA28/16 substrate. BSA was used as a negative control. (B) The effect of ED on RNA helicase activity of the full-length nsP2 (nsP2FL) protein was studied using the Alexa488-dsRNA28/16 substrate. The representative gel pictures of each experiment are shown in the top panel. The relative band intensities, quantified using ImageJ software, are provided in the bottom panels. The student’s t-test was used to calculate the *p* values: \**p*<0.05, \*\**p*<0.01, \*\*\**p*<0.001, \*\*\*\**p*<0.001, ns = not significant.

### ED inhibits CHIKV uptake

The other events that can affect the virus replication early during the CHIKV life cycle are the virus binding to the host cells and its subsequent uptake. CHIKV binding to ERMS cells was not affected at 0.1 µM ED concentration at which the virus RNA and protein synthesis, and viral replication was inhibited (Fig. 10A). However, virus uptake showed ∼30% inhibition at this concentration of ED (Fig. 10B). The inhibition of the virus binding and uptake at higher ED concentrations may be related to the inhibition of the host protein synthesis. The virucidal activity of ED was also ruled out when CHIKV titers were not affected when the virus was incubated with 20 µM ED (Fig. 10C).

**Figure 10.**
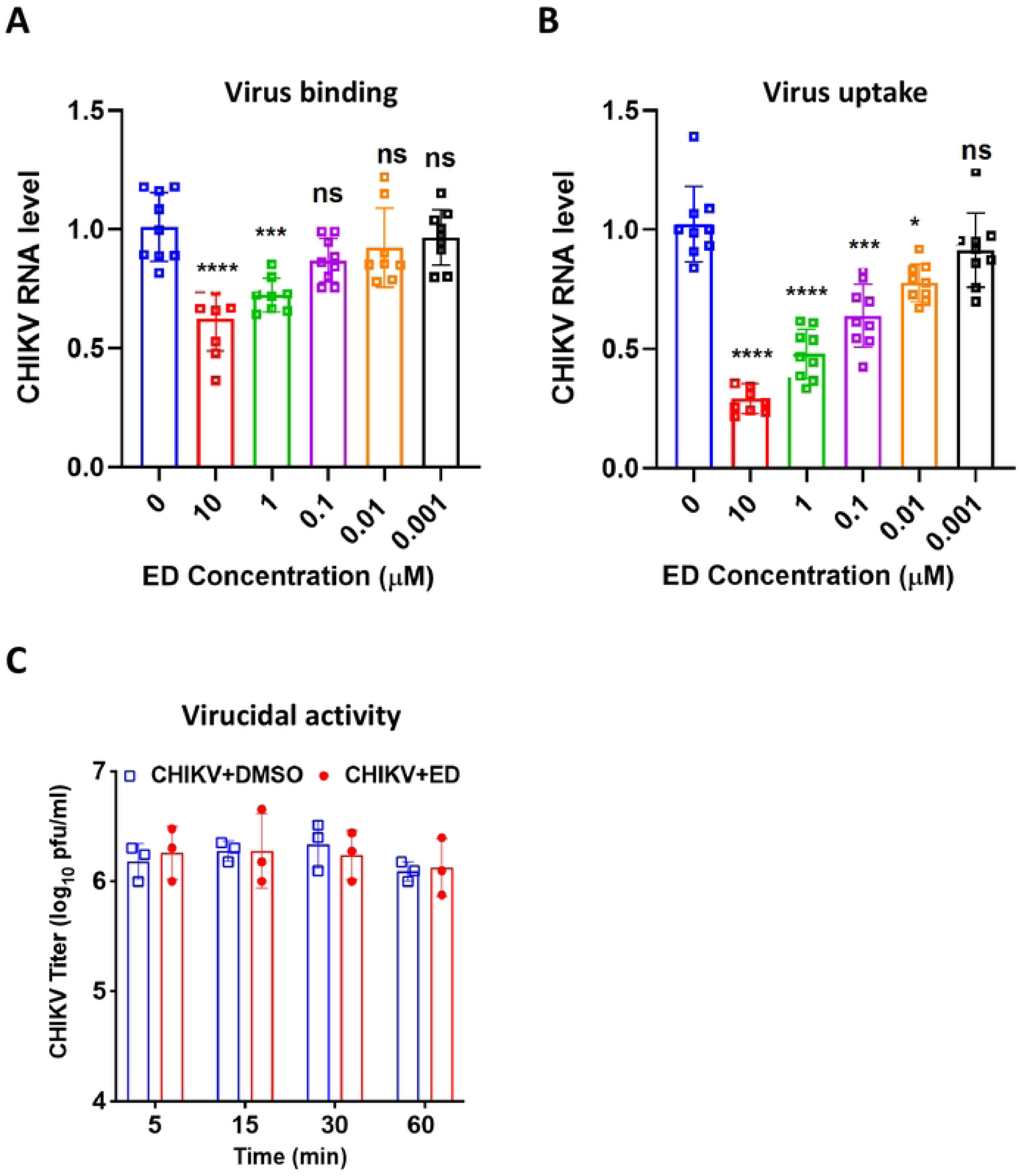
CHIKV binding and uptake in the host cells in the presence of ED. (A) ERMS cells were incubated for 1 h with different concentrations of ED at 37 °C, followed by the removal of the drug-containing culture medium. The cells were then incubated for 1 h with CHIKV (1 MOI) at 4 °C. Followed by a PBS wash, the cells were harvested for RNA isolation. The relative CHIKV RNA levels are shown, where *Gapdh* was used as the internal control. The viral RNA levels in the control (DMSO-treated) cells were compared with those treated with ED. (B) ERMS cells were incubated for 1 h with different concentrations of ED at 37 °C, followed by the removal of the drug-containing culture medium. The cells were then incubated for 1 h with CHIKV (1 MOI) at 4 °C, followed by incubation at 37 °C for 1 h. Following a PBS wash, the cells were incubated with trypsin to remove the extracellular virus, and the total RNA was isolated. The relative CHIKV RNA levels are shown, where *Gapdh* was used as the internal control. The viral RNA levels in the control (DMSO-treated) cells were compared with those treated with ED. The student’s t-test was used to calculate the p values: *p<0.05, **p<0.01, ***p<0.001, \*\*\*\**p*<0.0001, ns = not significant. (C) For the virucidal assay, CHIKV in DMEM was incubated with 20 µM ED or DMSO (control) for 1 h at 37 °C. The virus titers were then determined by plaque assay. The viral titers in the control were compared with those in the presence of ED by Student’s t-test; no significant differences in the viral titers were seen.

### ED inhibits the replication of different alphaviruses

The nsP2 amino acid sequence of alphaviruses Ross River virus (RRV) and Sindbis virus (SINV) showed about 69% and 58% sequence homology with the nsP2 protein of CHIKV (Fig. S1). Out of the nine amino acids of nsP2 helicase that lined ED within the 4.0 Å space in our *in silico* studies, eight were conserved in RRV, and seven in SINV (Fig. S1). We studied the effect of ED on the replication of these two viruses (Fig. 11). ED significantly reduced the viral RNA and viral titers in RRV- or SINV-infected ERMS cells at concentrations of 0.1 µM and as low as 0.001 µM.

**Figure 11.**
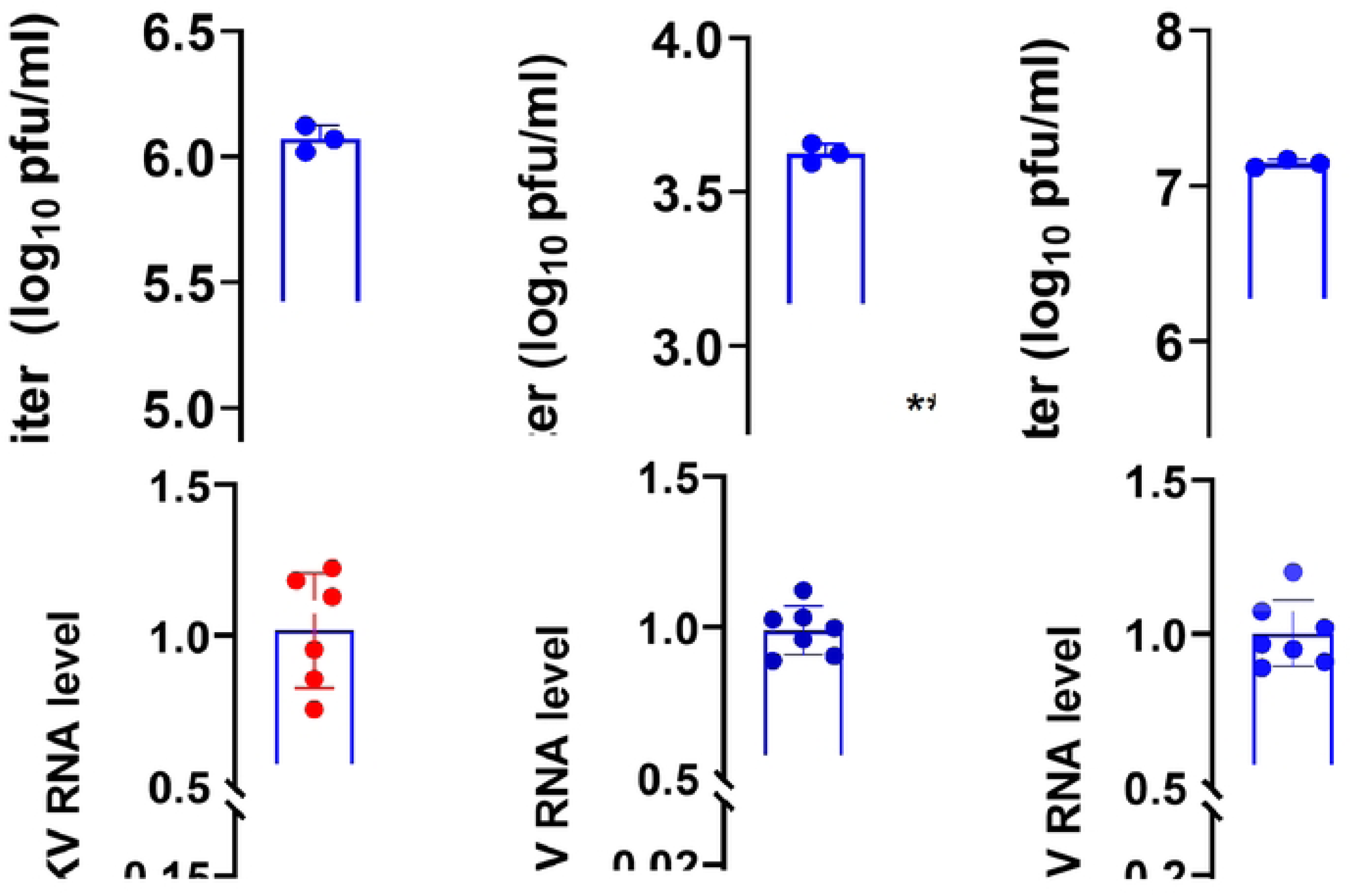
ED inhibits the replication of SINV and RRV. ERMS cells were infected with CHIKV, RRV or SINV (1 MOI) and incubated with different concentrations of ED. The cells and culture supernatants were harvested at 6 h pi for RNA extraction and viral titer determination, respectively. The viral titers determined by plaque assay are shown in the top panel. The relative viral RNA levels are shown in the bottom panel, where *Gapdh* was used as the internal control. The student’s t-test was used to calculate the *p* values: \**p*<0.05, \*\**p*<0.01, \*\*\**p*<0.001, \*\*\*\**p*<0.0001.

## DISCUSSION

A number of antiviral molecules have been identified against CHIKV using *in vitro* and *in silico* methods. However, these are still in the experimental phase and have not been taken through the drug development pathway, which is long, expensive, and tedious. Drug repurposing, or repositioning, is the process of finding new therapeutic uses for existing drugs that were originally developed for different indications. This approach is particularly advantageous for antiviral drug development because it leverages known safety profiles, manufacturing data, and pharmacokinetics of approved or investigational drugs, reducing the time and cost involved in bringing a new antiviral therapy to market. We have screened the Spectrum Collection of 2560 compounds that includes all the compounds in the US and International Drug Collections and have identified ED as a potent antiviral molecule against CHIKV in the cell culture and the mouse model of disease. The molecule with an IC_50_ of 10 nM showed a high selectivity index (SI) of 967, demonstrating its high potential for use as a CHIKV antiviral.

ED is the dichloride salt of emetine, a natural alkaloid derived from the roots of *Psychotria ipecacuanha* (ipecac), recognised for its emetic and antiprotozoal properties (46). Emetine has a long history of use, primarily as an antiemetic to induce vomiting, and antiprotozoal drug against *Entamoeba hystolytica* (47) until a safer metronidazole became available. The emetine-induced non-permanent cardiotoxicity was unequivocally associated with the high-dose (60 mg/day for 10 days) required to achieve a minimum inhibitory concentration of 25 µM against *Entamoeba hystolytica*. However, the toxicity was more commonly observed when treatment exceeded 5 days. No cardiovascular side effects were reported when emetine was used for various indications at a low dose of 20 mg/day (44).

In the past, emetine has been used to treat Spanish Influenza (44), Singles (48,49), adenoviral kerato-conjunctivitis (50), and hepatitis (51,52). In recent years, emetine has garnered attention as a potential therapeutic agent for viral infections, showing potent antiviral activity at IC_50_ values in sub-micromolar or low nanomolar ranges (44). A phase 2/3 trial is ongoing to determine emetine’s efficacy and safety for the treatment of symptomatic COVID-19 in a randomised clinical trial (ClinicalTrials.gov ID NCT05889793) at Johns Hopkins University, USA.

Emetine has demonstrated antiviral potential against a wide range of viruses, including RNA and DNA viruses. For instance, it was shown to inhibit the replication of the RNA genome containing Zika and Ebola viruses by inhibiting the replication and virus entry (53). Emetine inhibited Echovirus 1, Human metapneumovirus and Rift Valley fever virus (54), and coronaviruses such as hCoV-NL63, hCoV-OC43, MERS-CoV (55), and SARS-CoV-2 in cell culture (56–60). Besides, emetine exerted potent antiviral activity against Foot-and-mouth disease virus as a replicase inhibitor and direct virucide (60). DNA viruses, such as Herpes simplex virus type 2 (54) and Human cytomegalovirus (CMV), were inhibited by emetine using a specific host-dependent anti-CMV mechanism (61). The efficacy of emetine has also been demonstrated in preclinical studies in animal models against CMV (61), Zika virus (52), and Enterovirus-71(43).

CHIKV replication in skeletal muscle cells is required for disease development, and replication in skeletal muscle cells is a critical mediator of CHIKV pathogenesis (62). Mounting evidence has implicated skeletal muscle as an important site in CHIKV disease development (62–65). Therefore, in addition to the commonly used BHK cells, we studied emetine-mediated CHIKV inhibition in ERMS cells. These cells of human origin resemble undifferentiated muscle cells (myoblasts) and display characteristics of skeletal muscle lineage, like the expression of muscle-specific markers myogenin and desmin (67).

Emetine inhibited CHIKV replication in ERMS cells in the nanomolar range with an IC_50_ of 10 nM, and a complete inhibition of virus replication was seen at a 50 nM concentration. Importantly, in the mouse model of CHIKV infection, ED showed robust antiviral properties, where ED-treated CHIKV-infected mice showed lower viremia and did not develop characteristic footpad edema. The weight loss associated with CHIKV infection was also reversed in the ED-treated animals. A clear dose-effect was observed on clinical symptoms of viremia and footpad edema in CHIKV-infected mice treated with graded doses of ED. Mice that were treated with ED as late as 48 h after CHIKV exposure showed significantly reduced viremia and absence of clinical symptoms of footpad edema. ED was effective in mice at a subcutaneous dose of 6 mg/kg/day, which translates to a 0.49 mg/kg/day human dose. This is significantly lower than the 1 mg/kg/day dose used in humans to treat amoebiasis, which was causing some adverse effects (44). It may be noted that no cardiovascular side effects were reported when emetine was used for different indications at a low dose of 20 mg/day (44).

The ED-mediated inhibition of CHIKV replication, as studied by the kinetics of viral RNA synthesis and the extracellular virus titers in the infected cells, showed that ED inhibited the early stage/s of CHIKV replication. The significantly reduced CHIKV RNA levels in the virus-infected cells as early as 3 h pi indicate an inhibition of the virus binding or uptake step. While virus binding was not affected, CHIKV uptake was reduced in ERMS cells in the presence of ED. We ruled out any virucidal effect that ED might have on CHIKV virions by directly incubating the virus with ED. Previously, ED was shown to inhibit the entry of Ebola pseudovirus in HeLa cells (53), Hantaan virus in VeroE6 cells (68) SARS-CoV-2 in Vero cells(59), and Enterovirus-A71 in rhabdomyosarcoma cells (43). Although the mechanistic details were not studied.

The multifunctional nsP2 is one of the important CHIKV non-structural proteins, which has a protease domain containing the activity required for producing the mature non-structural proteins, and the helicase domain is essential for unwinding RNA to facilitate the viral RNA replication and transcription. Our data showed that ED strongly inhibited the replication of CHIKV RNA. Interestingly, our *in silico* studies predicted ED binding to the helicase domain of nsP2 with high affinity. This observation was experimentally validated using the ITC and MST methods, confirming the binding of ED with the helicase domain of nsP2 with a K_d_ value of 2.56 µM and 3.06 µM, respectively. Further, a mutation of the *in silico*-predicted ED-interacting nsP2 residue caused a significant reduction in the binding affinity of nsP2 with ED; the K_d_ value for the mutant nsP2 was 29.40 µM. Importantly, ED binding to the nsP2 protein inhibited its helicase activity *in vitro* in a dose-dependent manner. Our data thus clearly establish that ED suppresses CHIKV replication by binding to its nsP2 protein, inhibiting its helicase activity. Since we have not studied other CHIKV proteins, their role in the ED-mediated anti-CHIKV activity cannot be ruled out.

Emetine has been shown to exhibit its antiviral effects primarily by interfering with viral replication processes. One of its key mechanisms involves the inhibition of protein synthesis in host cells, which indirectly prevents the translation of viral proteins. Emetine achieves this by binding to the 40S ribosomal subunit, disrupting the elongation phase of translation (69). Additionally, emetine has been shown to disrupt the interaction between viral RNA and host cell translation machinery, specifically eukaryotic initiation factor 4E (eIF4E), which is critical for viral mRNA translation (70). The low IC_50_ of 10 nM and a complete inhibition of CHIKV replication at 50 nM ED suggest that the protein synthesis inhibition may have no direct role in the ED-mediated CHIKV inhibition in ERMS cells, since host cell protein synthesis was largely unaffected at 100 nM ED concentration.

The arthritogenic symptoms observed during the acute phase of CHIKV infection are largely due to the host’s pro-inflammatory responses, with elevated levels of cytokines and chemokines playing pivotal roles in the development of joint inflammation and pain (7,71). Emetine inhibits the activation of NF-κB, a transcription factor involved in inflammatory responses (72,73). Thus, modulation of host cell signalling pathways may also contribute to emetine-mediated inhibition of CHIKV replication and suppression of the footpad edema symptoms seen in the mouse model of CHIKV infection. Therefore, the role of NF-κB-mediated ED action in controlling CHIKV replication in the cells, and particularly in the animal model, needs investigation.

The robust anti-CHIKV activity in the cell culture and the therapeutic action of ED in the animal model, especially at a dose smaller than the prescribed human dose, establishes its antiviral potential. Additionally, emetine’s anti-inflammatory properties can mitigate the cytokine storm associated with CHIKV infection (74). An emetine concentration of 25 µM is required to inhibit *Entamoeba histolytica* growth *in vitro* (75), whereas near-complete inhibition of CHIKV replication was seen at a concentration of 50 nM in the cell culture. These concentrations are around 500-fold lower than the emetine dose required to kill the amoeba. These data warrant further studies on the use of emetine in treating CHIKV patients at doses lower than those used to treat amoebiasis. This will determine the feasibility of repurposing emetine as a viable CHIKV antiviral agent.

## FIGURE LEGENDS

**Figure S1. Conservation of the nsP2 sequence among CHIKV, SINV and RRV.**

The amino acid sequence of the CHIKV nsP2 protein was aligned with the SINV and RRV nsP2 proteins using the UniProt Align tool. The amino acids lining within the 4.0 Å space of the nsP2 interaction with ED, as identified *in silico*, are shown in red coloured boxes.

## Data Sharing Statement

All relevant data are within the manuscript and its supporting information files.

## Acknowledgments

The Small Animal Facility supported by the DBT grant no. BT/PR5480/INF/158/2012 is acknowledged. We also acknowledge the facilities and staff of the Advanced Technology Platform Centre (ATPC).

## Author contributions

**Conceptualisation:** Sudhanshu Vrati

**Data curation:** Anshula Sharma, Sudhanshu Vrati

**Formal analysis:** Anshula Sharma, Deepti Jain, Sudhanshu Vrati

**Funding acquisition:** Sudhanshu Vrati, Deepti Jain

**Investigation:** Anshula Sharma, Chandru Subramani, Shouri KA, Brohmomoy Basu, Ghanshyam Sharma, Archana Rout, Abhay Deep Pandey, Devendra Sharma

**Methodology:** Deepti Jain, Abhay Deep Pandey

**Project administration:** Sudhanshu Vrati

**Resources:** Deepti Jain

**Software:** Abhay Deep Pandey

**Supervision:** Deepti Jain, Sudhanshu Vrati

**Validation:** Anshula Sharma, Chandru Subramani, Deepti Jain

**Visualisation:** Anshula Sharma, Ghanshyam Sharma

**Writing – original draft:** Anshula Sharma

**Writing – review & editing**: Anshula Sharma, Abhay Deep Pandey, Deepti Jain, Sudhanshu Vrati

## References

1. Zeller H, Van Bortel W, Sudre B. Chikungunya: Its History in Africa and Asia and Its Spread to New Regions in 2013-2014. J Infect Dis. 2016;214(suppl 5):S436–40. Available from: https://pubmed.ncbi.nlm.nih.gov/27920169/.

2. Constant LEC, Rajsfus BF, Carneiro PH, Sisnande T, Mohana-Borges R, Allonso D. Overview on Chikungunya Virus Infection: From Epidemiology to State-of-the-Art Experimental Models. Front Microbiol. 2021;12:744164. Available from: https://pubmed.ncbi.nlm.nih.gov/34675908/.

3. Chikungunya worldwide overview [cited 2025 May 23]. Available from: https://www.ecdc.europa.eu/en/chikungunya-monthly.

4. Burt FJ, Chen W, Miner JJ, Lenschow DJ, Merits A, Schnettler E, et al. Chikungunya virus: an update on the biology and pathogenesis of this emerging pathogen. Lancet Infect Dis. 2017;17(4):e107–17. Available from: https://pubmed.ncbi.nlm.nih.gov/28159534/.

5. Arif M, Tauran P, Kosasih H, Pelupessy NM, Sennang N, Mubin RH, et al. Chikungunya in Indonesia: Epidemiology and diagnostic challenges. PLoS Negl Trop Dis. 2020;14(6):1–18. Available from: https://pubmed.ncbi.nlm.nih.gov/32479497/.

6. Hakim MS, Annisa L, Aman AT. The evolution of chikungunya virus circulating in Indonesia: Sequence analysis of the orf2 gene encoding the viral structural proteins. Int Microbiol. 2023;26(4):781–90. Available from: https://pubmed.ncbi.nlm.nih.gov/36774411/.

7. Pathak H, Mohan MC, Ravindran V. Chikungunya arthritis. Clin Med (Lond). 2019;19(5):381–5. Available from: https://pubmed.ncbi.nlm.nih.gov/31530685/.

8. Khan AH, Morita K, del Carmen Parquet M, Hasebe F, Mathenge EGM, Igarashi A. Complete nucleotide sequence of chikungunya virus and evidence for an internal polyadenylation site. J Gen Virol. 2002;83(Pt 12):3075–84. Available from: https://pubmed.ncbi.nlm.nih.gov/12466484/.

9. Frolov I, Frolova EI. Molecular Virology of Chikungunya Virus. Curr Top Microbiol Immunol. 2022;435:1–31. Available from: https://pubmed.ncbi.nlm.nih.gov/30599050/.

10. Bartholomeeusen K, Daniel M, LaBeaud DA, Gasque P, Peeling RW, Stephenson KE, et al. Chikungunya fever. Nat Rev Dis Primers. 2023;9(1). Available from: https://pubmed.ncbi.nlm.nih.gov/37024497/.

11. Delang L, Li C, Tas A, Quérat G, Albulescu IC, De Burghgraeve T, et al. The viral capping enzyme nsP1: a novel target for the inhibition of chikungunya virus infection. Sci Rep. 2016;6:31819. Available from: https://pubmed.ncbi.nlm.nih.gov/27545976/.

12. Kovacikova K, Morren BM, Tas A, Albulescu IC, Van Rijswijk R, Jarhad DB, et al. 6’-β-Fluoro-Homoaristeromycin and 6’-Fluoro-Homoneplanocin A Are Potent Inhibitors of Chikungunya Virus Replication through Their Direct Effect on Viral Nonstructural Protein 1. Antimicrob Agents Chemother. 2020;64(4):e02532–19. Available from: https://pubmed.ncbi.nlm.nih.gov/31964798/.

13. Dansana J, Purohit P, Panda M, Meher BR. Recent advances in phytocompounds as potential Chikungunya virus non-structural protein 2 protease antagonists: A systematic review. Phytomedicine. 2025;136:156359. Available from: https://pubmed.ncbi.nlm.nih.gov/39756312/.

14. Setyawati I, Setiawan AG, Nemchinova M, Vidilaseris K. The potential inhibitory mechanism of EGCG against the Chikungunya virus targeting non-structural protein 2 through molecular dynamics simulation. Sci Rep. 2024;14(1):29797. Available from: https://pubmed.ncbi.nlm.nih.gov/39616212/.

15. Sharma KB, Subramani C, Ganesh K, Sharma A, Basu B, Balyan S, et al. Withaferin A inhibits Chikungunya virus nsP2 protease and shows antiviral activity in the cell culture and mouse model of virus infection. PLoS Pathog. 2024;20(12):e1012816. Available from: https://pubmed.ncbi.nlm.nih.gov/39775571/.

16. Chaudhary M, Kumar A, Bala Sharma K, Vrati S, Sehgal D. In silico identification of chikungunya virus replication inhibitor validated using biochemical and cell-based approaches. FEBS J. 2024;291(12):2656–73. Available from: https://pubmed.ncbi.nlm.nih.gov/38303163/.

17. Battisti V, Urban E, Langer T. Antivirals against the Chikungunya Virus. Viruses. 2021;13(7):1307. Available from: https://pubmed.ncbi.nlm.nih.gov/34372513/.

18. Ghildiyal R, Gabrani R. Computational approach to decipher cellular interactors and drug targets during co-infection of SARS-CoV-2, Dengue, and Chikungunya virus. Virus disease. 2021;32(1):55–64. Available from: https://pubmed.ncbi.nlm.nih.gov/33723515/.

19. Henss L, Scholz T, Grünweller A, Schnierle BS. Silvestrol Inhibits Chikungunya Virus Replication. Viruses. 2018;10(11):592. Available from: https://pubmed.ncbi.nlm.nih.gov/30380742/.

20. Langsjoen RM, Auguste AJ, Rossi SL, Roundy CM, Penate HN, Kastis M, et al. Host oxidative folding pathways offer novel anti-chikungunya virus drug targets with broad spectrum potential. Antiviral Res. 2017;143:246–51. Available from: https://pubmed.ncbi.nlm.nih.gov/28461071/.

21. Rathore APS, Haystead T, Das PK, Merits A, Ng ML, Vasudevan SG. Chikungunya virus nsP3 & nsP4 interacts with HSP-90 to promote virus replication: HSP-90 inhibitors reduce CHIKV infection and inflammation in vivo. Antiviral Res. 2014;103(1):7–16. Available from: https://pubmed.ncbi.nlm.nih.gov/24388965/.

22. Castro EF, Álvarez DE. New Highly Selective Antivirals for Chikungunya Virus identified from the Screening of a Drug-Like Compound Library. Curr Microbiol. 2024;81(10). Available from: https://pubmed.ncbi.nlm.nih.gov/39227496/

23. Metibemu DS, Adeyinka OS, Falode J, Hampton T, Crown O, Ojobor JC, et al. Inhibitor of the non-structural protein 2 protease shows promising efficacy in mouse models of chikungunya. Eur J Med Chem. 2024;278:116808. Available from: https://pubmed.ncbi.nlm.nih.gov/39236495/.

24. Liu Y, Xu M, Xia B, Qiao Z, He Y, Liu Y, et al. Nifuroxazide Prevents Chikungunya Virus Infection Both In Vitro and In Vivo via Suppressing Viral Replication. Viruses. 2024;16(8):1322. Available from: https://pubmed.ncbi.nlm.nih.gov/39205296/.

25. Loaiza-Cano V, Hernández-Mira E, Pastrana-Restrepo M, Galeano E, Pardo-Rodriguez D, Martinez-Gutierrez M. The Mechanism of Action of L-Tyrosine Derivatives against Chikungunya Virus Infection In Vitro Depends on Structural Changes. Int J Mol Sci. 2024;25(14):7972. Available from: https://pubmed.ncbi.nlm.nih.gov/39063216/.

26. Agrawal T, Siddqui G, Dahiya R, Patidar A, Madan U, Das S, et al. Inhibition of early RNA replication in Chikungunya and Dengue virus by lycorine: In vitro and in silico studies. Biochem Biophys Res Commun. 2024;730:150393. Available from: https://pubmed.ncbi.nlm.nih.gov/39003865/.

27. Hucke FIL, Bugert JJ. Current and Promising Antivirals Against Chikungunya Virus. Front Public Health. 2020;8:618624. Available from: https://pubmed.ncbi.nlm.nih.gov/33384981/.

28. Kovacikova K, van Hemert MJ. Small-Molecule Inhibitors of Chikungunya Virus: Mechanisms of Action and Antiviral Drug Resistance. Antimicrob Agents Chemother. 2020;64(12):e01788–20. Available from: https://pubmed.ncbi.nlm.nih.gov/32928738/.

29. Rabelo VW, Sanchez-Nuñez ML, Corrêa-Amorim LS, Kuhn RJ, Abreu PA, Paixão ICNP. In Silico Drug Repurposing Uncovered the Antiviral Potential of the Antiparasitic Drug Oxibendazole Against the Chikungunya Virus. ACS Omega. 2024;9(25):27632–42. Available from: https://pubmed.ncbi.nlm.nih.gov/38947813/.

30. Souza BG de, Choudhary S, Vilela GG, Passos GFS, Costa CACB, Freitas JD de, et al. Design, synthesis, antiviral evaluation, and In silico studies of acrylamides targeting nsP2 from Chikungunya virus. Eur J Med Chem. 2023;258:115572. Available from: https://pubmed.ncbi.nlm.nih.gov/37364511/.

31. Verma J, Hasan A, Sunil S, Subbarao N. In silico identification and in vitro antiviral validation of potential inhibitors against Chikungunya virus. J Comput Aided Mol Des. 2022;36(7):521–36. Available from: https://pubmed.ncbi.nlm.nih.gov/35789450/.

32. Thomas N, Patil P, Sharma A, Kumar S, Singh VK, Alagarasu K, et al. Studies on the antiviral activity of chebulinic acid against dengue and chikungunya viruses and in silico investigation of its mechanism of inhibition. Sci Rep. 2022;12(1):10397. Available from: https://pubmed.ncbi.nlm.nih.gov/35729191/.

33. Chaudhary M, Sehgal D. In silico identification of natural antiviral compounds as a potential inhibitor of chikungunya virus non-structural protein 3 macrodomain. J Biomol Struct Dyn. 2022;40(22):11560–70. Available from: https://pubmed.ncbi.nlm.nih.gov/34355667/.

34. Sangeetha K, Purushothaman I, Rajarajan S. Spectral characterisation, antiviral activities, in silico ADMET and molecular docking of the compounds isolated from Tectona grandis to chikungunya virus. Biomed Pharmacother. 2017;87:302–10. Available from: https://pubmed.ncbi.nlm.nih.gov/28063412/.

35. Tssetsarkin K, Higgs S, McGee CE, De Lamballerie X, Charrel RN, Vanlandingham DL. Infectious clones of Chikungunya virus (La Réunion isolate) for vector competence studies. Vector Borne Zoonotic Dis. 2006;6(4):325–37. Available from: https://pubmed.ncbi.nlm.nih.gov/17187566/.

36. Vashist S, Urena L, Goodfellow I. Development of a strand specific real-time RT-qPCR assay for the detection and quantitation of murine norovirus RNA. J Virol Methods. 2012;184(1–2):69–76. Available from: https://pubmed.ncbi.nlm.nih.gov/22626565/.

37. Sidorenko VS, Cohen I, Dorjee K, Minetti CA, Remeta DP, Gao J, et al. Mechanisms of antiviral action and toxicities of ipecac alkaloids: Emetine and dehydroemetine exhibit anti-coronaviral activities at non-cardiotoxic concentrations. Virus Res. 2024;341:199322. Available from: https://pubmed.ncbi.nlm.nih.gov/38228190/.

38. Law YS, Wang S, Tan YB, Shih O, Utt A, Goh WY, et al. Interdomain Flexibility of Chikungunya Virus nsP2 Helicase-Protease Differentially Influences Viral RNA Replication and Infectivity. J Virol. 2021;95(6):e01470–20. Available from: https://pmc.ncbi.nlm.nih.gov/articles/PMC8094934/.

39. Das PK, Merits A, Lulla A. Functional cross-talk between distant domains of chikungunya virus non-structural protein 2 is decisive for its RNA-modulating activity. J Biol Chem. 2014;289(9):5635–53. Available from: https://pubmed.ncbi.nlm.nih.gov/24407286/.

40. Pastorino BAM, Peyrefitte CN, Almeras L, Grandadam M, Rolland D, Tolou HJ, et al. Expression and biochemical characterization of nsP2 cysteine protease of Chikungunya virus. Virus Res. 2008;131(2):293–8. Available from: https://pubmed.ncbi.nlm.nih.gov/17961784/.

41. Law YS, Utt A, Tan YB, Zheng J, Wang S, Chen MW, et al. Structural insights into RNA recognition by the Chikungunya virus nsP2 helicase. Proc Natl Acad Sci U S A. 2019;116(19):9558–67. Available from: https://pubmed.ncbi.nlm.nih.gov/31000599/.

42. Yousefi H, Mashouri L, Okpechi SC, Alahari N, Alahari SK. Repurposing existing drugs for the treatment of COVID-19/SARS-CoV-2 infection: A review describing drug mechanisms of action. Biochem Pharmacol. 2021;183:114296. Available from: https://pubmed.ncbi.nlm.nih.gov/33191206/.

43. Tang Q, Li S, Du L, Chen S, Gao J, Cai Y, et al. Emetine protects mice from enterovirus infection by inhibiting viral translation. Antiviral Res. 2020;173:104650. Available from: https://www.sciencedirect.com/science/article/pii/S0166354219304954.

44. Bleasel MD, Peterson GM. Emetine Is Not Ipecac: Considerations for Its Use as Treatment for SARS-CoV2. Pharmaceuticals. 2020;13(12):428. Available from: https://www.mdpi.com/1424-8247/13/12/428/htm.

45. Kasabe B, Ahire G, Patil P, Punekar M, Davuluri KS, Kakade M, et al. Drug repurposing approach against chikungunya virus: an in vitro and in silico study. Front Cell Infect Microbiol. 2023;13:1132538. Available from: https://pubmed.ncbi.nlm.nih.gov/37180434/.

46. Lee MR. Ipecacuanha: the South American vomiting root. J R Coll Physicians Edinb. 2008;38(4):355–60. Available from: https://pubmed.ncbi.nlm.nih.gov/19227966/.

47. Scragg JN, Powell SJ. Emetine hydrochloride and chloroquine in the treatment of children with amoebic liver abscess. Arch Dis Child. 1966;41(219):549–50. Available from: https://pubmed.ncbi.nlm.nih.gov/5957731/.

48. Viegas LB, Viegas LC. Shingles and emetine hydrochloride. Ann Dermatol Syphiligr (Paris). 1957;84(4):400–405. Available from: https://pubmed.ncbi.nlm.nih.gov/13458910/.

49. Annamalai R. Emetine hydrochloride in the treatment of herpes zoster. Indian J Dermatol. 1965;10:61. Available from: https://pubmed.ncbi.nlm.nih.gov/14266439/.

50. Hanisch J. New treatment of keratoconjunctivitis using epidemica emetine. Szemeszet. 1963;99:29–33.

51. Del Puerto BM, Tato JC, Koltan A, Bures OM, De Chieri PR, Garcia A, Escaray TI, Lorenzo B. Viral hepatitis in children and special reference to its treatment with emetine. Prensa Medica Argentina. 1968;55(18):818–34. Available from: https://pubmed.ncbi.nlm.nih.gov/5747742/.

52. Fusillo A. Effect of small doses of emetine in the therapy of virus diseases. New specific therapeutic method in viral hepatitis. Minerva medica. 1973;64(3):129–32. Available from: https://pubmed.ncbi.nlm.nih.gov/4686977/.

53. Yang S, Xu M, Lee EM, Gorshkov K, Shiryaev SA, He S, et al. Emetine inhibits Zika and Ebola virus infections through two molecular mechanisms: inhibiting viral replication and decreasing viral entry. Cell Discov. 2018;4(1):31. Available from: https://pubmed.ncbi.nlm.nih.gov/29872540/.

54. Andersen PI, Krpina K, Ianevski A, Shtaida N, Jo E, Yang J, et al. Novel Antiviral Activities of Obatoclax, Emetine, Niclosamide, Brequinar, and Homoharringtonine. Viruses. 2019;11(10):964. Available from: https://pubmed.ncbi.nlm.nih.gov/31635418/.

55. Shen L, Niu J, Wang C, Huang B, Wang W, Zhu N, et al. High-Throughput Screening and Identification of Potent Broad-Spectrum Inhibitors of Coronaviruses. J Virol. 2019;93(12):10–128. Available from: https://pubmed.ncbi.nlm.nih.gov/30918074/.

56. Choy KT, Wong AYL, Kaewpreedee P, Sia SF, Chen D, Hui KPY, et al. Remdesivir, lopinavir, emetine, and homoharringtonine inhibit SARS-CoV-2 replication in vitro. Antiviral Res. 2020;178:104786. Available from: https://pubmed.ncbi.nlm.nih.gov/32251767/.

57. Bojkova D, Klann K, Koch B, Widera M, Krause D, Ciesek S, et al. Proteomics of SARS-CoV-2-infected host cells reveals therapy targets. Nature. 2020;583(7816):469–72. Available from: https://pubmed.ncbi.nlm.nih.gov/32408336/.

58. Jan JT, Cheng TJR, Juang YP, Ma HH, Wu YT, Yang W Bin, et al. Identification of existing pharmaceuticals and herbal medicines as inhibitors of SARS-CoV-2 infection. Proc Natl Acad Sci U S A. 2021;118(5):e2021579118. Available from: https://pubmed.ncbi.nlm.nih.gov/33452205/.

59. Wang A, Sun Y, Liu Q, Wu H, Liu J, He J, et al. Low dose of emetine as potential anti-SARS-CoV-2 virus therapy: preclinical in vitro inhibition and in vivo pharmacokinetic evidences. Molecular biomedicine. 2020;1(1);1–9. Available from: https://pubmed.ncbi.nlm.nih.gov/34765997/.

60. Pantanam A, Mana N, Semkum P, Lueangaramkul V, Phecharat N, Lekcharoensuk P, et al. Dual effects of ipecac alkaloids with potent antiviral activity against foot-and-mouth disease virus as replicase inhibitors and direct virucides. Int J Vet Sci Med. 2024;12(1):134–47. Available from: https://pubmed.ncbi.nlm.nih.gov/39359867/.

61. Mukhopadhyay R, Roy S, Venkatadri R, Su YP, Ye W, Barnaeva E, et al. Efficacy and Mechanism of Action of Low Dose Emetine against Human Cytomegalovirus. PLoS Pathog. 2016;12(6):e1005717. Available from: https://pubmed.ncbi.nlm.nih.gov/27336364/.

62. Lentscher AJ, McCarthy MK, May NA, Davenport BJ, Montgomery SA, Raghunathan K, et al. Chikungunya virus replication in skeletal muscle cells is required for disease development. J Clin Invest. 2020;130(3):1466–78. Available from: https://pubmed.ncbi.nlm.nih.gov/31794434/.

63. Rohatgi A, Corbo JC, Monte K, Higgs S, Vanlandingham DL, Kardon G, et al. Infection of myofibers contributes to increased pathogenicity during infection with an epidemic strain of chikungunya virus. J Virol. 2014;88(5):2414–25. Available from: https://pubmed.ncbi.nlm.nih.gov/24335291/.

64. Hawman DW, Carpentier KS, Fox JM, May NA, Sanders W, Montgomery SA, et al. Mutations in the E2 Glycoprotein and the 3’ Untranslated Region Enhance Chikungunya Virus Virulence in Mice. J Virol. 2017;91(20). Available from: https://pubmed.ncbi.nlm.nih.gov/28747508/.

65. Nair S, Poddar S, Shimak RM, Diamond MS. Interferon Regulatory Factor 1 Protects against Chikungunya Virus-Induced Immunopathology by Restricting Infection in Muscle Cells. J Virol. 2017;91(22):10–128. Available from: https://pubmed.ncbi.nlm.nih.gov/28835505/.

66. Couderc T, Chrétien F, Schilte C, Disson O, Brigitte M, Guivel-Benhassine F, et al. A mouse model for Chikungunya: young age and inefficient type-I interferon signaling are risk factors for severe disease. PLoS Pathog. 2008;4(2):e29. Available from: https://pubmed.ncbi.nlm.nih.gov/18282093/.

67. Li RF, Gupta M, McCluggage WG, Ronnett BM. Embryonal rhabdomyosarcoma (botryoid type) of the uterine corpus and cervix in adult women: report of a case series and review of the literature. Am J Surg Pathol. 2013;37(3):344–55. Available from: https://pubmed.ncbi.nlm.nih.gov/23348207/.

68. Mayor J, Torriani G, Engler O, Rothenberger S. Identification of Novel Antiviral Compounds Targeting Entry of Hantaviruses. Viruses. 2021;13(4):685. Available from: https://pubmed.ncbi.nlm.nih.gov/33923413/.

69. Wong W, Bai XC, Brown A, Fernandez IS, Hanssen E, Condron M, et al. Cryo-EM structure of the Plasmodium falciparum 80S ribosome bound to the anti-protozoan drug emetine. Elife. 2014;3(3):e03080. Available from: https://pubmed.ncbi.nlm.nih.gov/24913268/.

70. Kumar R, Afsar M, Khandelwal N, Chander Y, Riyesh T, Dedar RK, et al. Emetine suppresses SARS-CoV-2 replication by inhibiting interaction of viral mRNA with eIF4E. Antiviral Res. 2021;189:105056. Available from: https://pubmed.ncbi.nlm.nih.gov/33711336/.

71. Wauquier N, Becquart P, Nkoghe D, Padilla C, Ndjoyi-Mbiguino A, Leroy EM. The acute phase of Chikungunya virus infection in humans is associated with strong innate immunity and T CD8 cell activation. J Infect Dis. 2011;204(1):115–23. Available from: https://pubmed.ncbi.nlm.nih.gov/21628665/.

72. Miller SC, Huang R, Sakamuru S, Shukla SJ, Attene-Ramos MS, Shinn P, et al. Identification of known drugs that act as inhibitors of NF-kappaB signaling and their mechanism of action. Biochem Pharmacol. 2010;79(9):1272–80. Available from: https://pubmed.ncbi.nlm.nih.gov/20067776/.

73. Silva SLR, Dias IRSB, Rodrigues ACB da C, Costa RGA, Oliveira M de S, Barbosa GA da C, et al. Emetine induces oxidative stress, cell differentiation and NF-κB inhibition, suppressing AML stem/progenitor cells. Cell Death Discov. 2024;10(1):201. Available from: https://pubmed.ncbi.nlm.nih.gov/38684672/.

74. Morel Z, Martínez T, Galeano F, Coronel J, Quintero L, Jimenez R, et al. Cytokine storm in Chikungunya: Can we call it multisystem inflammatory syndrome associated with Chikungunya? Reumatol Clin. 2024;20(4):223–5. Available from: https://pubmed.ncbi.nlm.nih.gov/38644032/.

75. Burchard GD, Mirelman D. Entamoeba histolytica: virulence potential and sensitivity to metronidazole and emetine of four isolates possessing nonpathogenic zymodemes. Exp Parasitol. 1988;66(2):231–42. Available from: https://pubmed.ncbi.nlm.nih.gov/2899517/.

